# Discovery of chromatin-based determinants of azacytidine and decitabine anti-cancer activity

**DOI:** 10.1101/2025.07.29.664810

**Authors:** Rishi V. Puram, Qiangzong Yin, YuhJong Liu, Justine C. Rutter, Daniel Bondeson, Maria C. Saberi, Lisa Miller, Michael Du, Khanh Nguyen, Donovan L. Batzli, Hilina B. Woldemichael, Christian Täger, Anna Goldstein, Ming Y. Chu, Qi Guo, D. R. Mani, Michael Naumann, Melissa M. Ronan, Matthew G. Rees, Blanche C. Ip, Mustafa Kocak, Mikolaj Slabicki, John G. Doench, Jennifer A. Roth, Steven A. Carr, Namrata D. Udeshi, Jingyi Wu, Todd R. Golub

## Abstract

The DNA-incorporating nucleoside analogs azacytidine (AZA) and decitabine (DEC) have clinical efficacy in blood cancers, yet the precise mechanism by which these agents kill cancer cells has remained unresolved – specifically, whether their anti-tumor activity arises from conventional DNA damage or DNA hypomethylation via DNA methyltransferase 1 (DNMT1) inhibition. This incomplete mechanistic understanding has limited their broader therapeutic application, particularly in solid tumors, where early clinical trials showed limited efficacy. Here, through the assessment of drug sensitivity in over 600 human cancer models and comparison to a non-DNA-damaging DNMT1 inhibitor (GSK-3685032), we establish DNA hypomethylation, rather than DNA damage, as the primary killing mechanism of AZA and DEC across diverse cancer types. In further support of an epigenetic killing mechanism, CRISPR drug modifier screens identified a core set of chromatin regulators, most notably the histone deubiquitinase USP48, as AZA and DEC protective factors. We show that USP48 is recruited to newly hypomethylated CpG islands and deubiquitinates non-canonical histones, establishing USP48 as a key molecular link between the two components of epigenetic gene regulation: DNA methylation and chromatin modification. Furthermore, loss of *USP48*, which occurs naturally through biallelic deletions in human cancers, sensitized both hematologic and solid tumors to DNMT1 inhibition *in vitro* and *in vivo*. Our findings elucidate the epigenetic mechanism of action of AZA and DEC and identify a homeostatic link between DNA methylation and chromatin state, revealing new therapeutic opportunities for DNMT1 inhibitors in solid tumors.

## INTRODUCTION

The anti-cancer agents 5-azacytidine (azacytidine; AZA) and 5-aza-2’-deoxycytidine (decitabine; DEC) have emerged as effective therapeutics for myelodysplastic syndromes (MDS) and acute myeloid leukemia (AML), largely through empiric trials^1–3^. AZA and DEC were first synthesized in 1960 as DNA-incorporating cytidine nucleoside analogs with a nitrogen in the fifth position of their pyrimidine ring. These compounds were initially developed as chemotherapeutic agents given their structural similarity to the DNA-damaging drug cytosine arabinoside (cytarabine; ARA-C)^4^. Although early studies with AZA and DEC revealed unexpected properties — including preserved efficacy in chemotherapy-resistant leukemias^5^ and the capacity to induce cellular transdifferentiation through DNA methyltransferase inhibition^6–10^ — the precise mechanism underlying their anti-neoplastic activity is widely debated^11–16^.

Systematic approaches to assess the molecular mechanism by which these drugs kill tumor cells have been lacking. AZA and DEC are incorporated into DNA during replication and form irreversible covalent complexes with DNA methyltransferase 1 (DNMT1), the enzyme responsible for maintaining DNA methylation patterns during DNA replication^17,18^. This triggers tumor cell killing through two potential mechanisms: conventional DNA damage from DNA-DNMT1 crosslinks and DNA hypomethylation from DNMT1 inhibition^19–21^. The relative contributions of these mechanisms to therapeutic activity are unclear. Furthermore, while AZA and DEC have demonstrated clinical utility in treating blood cancers, their potential efficacy in solid tumors remains incompletely explored^22^. Early clinical trials of the drugs in solid tumors showed minimal success, but these studies were limited by two crucial gaps: non-optimal dosing schedules and inability to predict which patients will respond due to uncertainty about the drug killing mechanism^23–25^.

Recent technological advances have created new opportunities to dissect these long-standing questions. First, the development of non-covalent DNMT1 inhibitors that do not incorporate into DNA enables the specific investigation of DNA hypomethylation effects in cancer cells without confounding DNA damage^26^. Second, genome-wide CRISPR screening now allows for the systematic identification of genes modulating drug response, and multiplexed cell barcoding methods can be used to measure drug sensitivity across hundreds of cancer models simultaneously^27,28^.

Here, we establish DNA hypomethylation as the primary tumor cell killing mechanism of AZA and DEC, while uncovering the core epigenetic circuitry that determines cellular response to DNA hypomethylation. Through systematic screening across hundreds of cancer models, we identify unexpected tumor types sensitive to DNMT1 inhibition and discover new chromatin-based factors, in particular the histone deubiquitinase (DUB) Ubiquitin Specific Protease 48 (USP48), that modulate drug response. Our findings reveal mechanistic principles governing the relationship between DNA methylation and chromatin regulation while expanding the therapeutic scope of DNMT1 inhibitors in cancer.

## RESULTS

### AZA and DEC exhibit DNMT1-dependent killing activity in diverse cancer types

To gain insight into the mechanism of action of AZA and DEC, we systematically profiled their anti-cancer activity across hundreds of human cancer cell line models using a high-throughput molecular barcoding method called PRISM (Profiling Relative Inhibition Simultaneously in Mixtures)^28^. We screened 620 DNA-barcoded cancer cell lines representing more than 20 tumor lineages and assessed cell viability by measuring relative barcode abundance following drug treatment for 6 days, compared to a DMSO control (Extended Data Fig. 1A). The cell killing patterns for AZA and DEC across the cancer models were highly correlated with each other (r = 0.74), confirming similar mechanisms of action despite the structural difference in their sugar backbones (Fig. 1A and Supplementary Table 1). This finding is consistent with the observation that the ribose sugar of AZA is converted to a deoxyribose form by ribonucleotide reductase^29^, thereby resulting in DNA incorporation and pharmacologic effects similar to DEC. Furthermore, this result indicates that RNA incorporation of AZA is not its primary anti-tumor mode of action.

**Figure 1:**
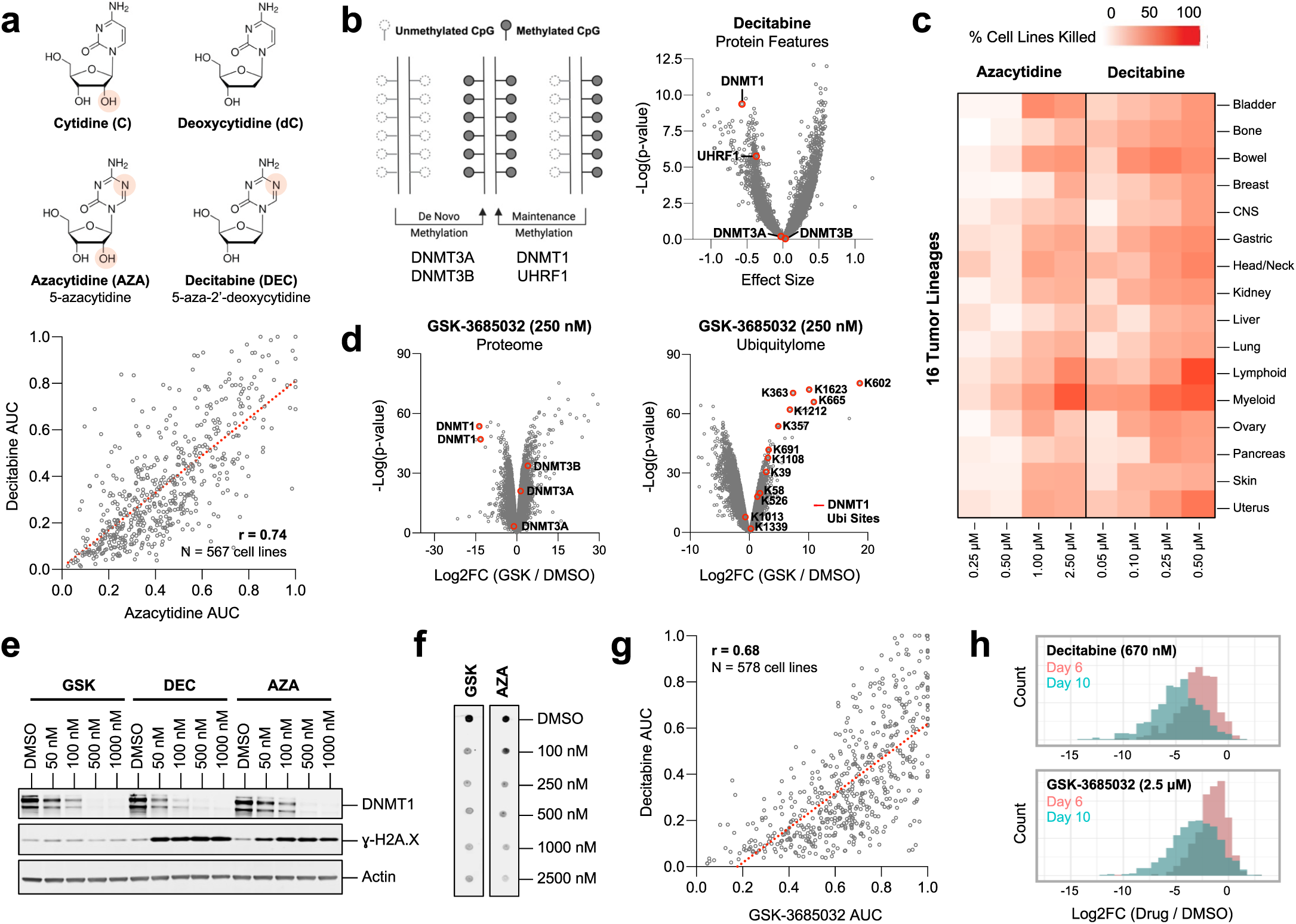
DNA hypomethylation via DNMT1 degradation is the principal killing mechanism of AZA and DEC in diverse cancers. **a.** Upper panel: Chemical structures of AZA and DEC, highlighting the shared nitrogen substitution in the fifth position of their pyrimidine rings and distinct sugar backbones. Lower panel: AUC correlation analysis of AZA and DEC across all cell lines in the PRISM screen. **b.** Left panel: The *de novo* methyltransferases DNMT3A and DNMT3B act on unmethylated DNA, whereas the maintenance methyltransferase DNMT1 and its binding partner UHRF1 act on hemimethylated DNA during DNA replication. Right panel: Analysis of protein expression features across the PRISM cell lines predictive of DEC sensitivity. **c.** Lineage analysis of AZA and DEC killing activity in the PRISM screen. Heatmap indicates the percentage of sensitive cell lines (Log2FC<-2) within each lineage for the lowest four doses of each drug. **d.** Comprehensive profiling of the proteome (left) and ubiquitylome (normalized to proteome) (right) in MV4;11 leukemia cells following treatment with 250 nM GSK-3685032 (n=3) for 48 hrs, as compared to DMSO (n=3). Each datapoint represents a protein group (see Methods). **e.** Western blot analysis of DNMT1 protein and γ-H2A.X levels in MOLM-13 cells following 48-hr treatment with AZA, DEC, or GSK-3685032. **f.** 5-methylcytosine DNA dot blot of MOLM-13 cells after 72-hr treatment with AZA or GSK-3685032. **g.** AUC correlation analysis of DEC and GSK-3685032 across the cancer cell lines in PRISM. **h.** Viability histograms for >500 cell lines treated with DEC (n=3) (upper panel) or GSK-3685032 (n=3) (lower panel) for 6 or 10 days in an extended timecourse PRISM screen.

To understand the mechanism responsible for tumor cell killing by AZA and DEC, we analyzed genomic features of each of the cell lines to identify predictive biomarkers of response. High protein expression of DNMT1 and its binding partner UHRF1^30^ – but not the *de novo* methyltransferases DNMT3A or DNMT3B^31^ – emerged as strong predictors of sensitivity to both AZA and DEC (Fig. 1B, Extended Data Fig. 1B, and Supplementary Table 2). This result validates DNMT1 as the primary molecular target of these drugs. Interestingly, *DNMT1* RNA expression (as opposed to protein levels) showed poor correlation with drug sensitivity, consistent with post-translational regulation of DNMT1 (Extended Data Fig. 1C and Supplementary Table 2).

Importantly, AZA and DEC potently killed not only leukemia cell lines but also subsets of solid tumor models, with notable efficacy in previously unreported contexts such as gastric, bladder, colon, ovarian, and head and neck cancers (Fig. 1C). Single cell line validation confirmed that both compounds induced killing in a variety of cancer lineages, suggesting that AZA and DEC should be explored further as solid tumor agents (Extended Data Fig. 1D-F).

### GSK-3685032 killing activity parallels AZA and DEC

The PRISM results suggested that cell killing by AZA and DEC was through a DNMT1-dependent mechanism. To more definitively test the hypothesis that the primary mode of killing by AZA and DEC is DNMT1 inhibition and DNA hypomethylation, we extended the PRISM analysis to a newly described compound, GSK-3685032, that inhibits DNMT1 through reversible, non-covalent binding without DNA incorporation^26^. Comprehensive proteomic profiling^32^ revealed that GSK-3685032 induced selective DNMT1 ubiquitination and degradation, while sparing DNMT3A and DNMT3B, confirming the compound’s specificity (Fig. 1D and Supplementary Table 3). Whereas AZA, DEC, and GSK-3685032 all induced dose-dependent DNMT1 protein degradation and global DNA hypomethylation, only AZA and DEC treatment resulted in increased levels of γ-H2A.X, a measure of DNA double-strand break formation (Fig. 1E-F and Extended Data Fig. 1G).

PRISM screening of 620 cancer cell lines revealed that DNA hypomethylation alone, induced by GSK-3685032, was sufficient to kill cells from multiple lineages (Extended Data Fig. 1H-I). Moreover, the cell killing pattern of GSK-3685032 was strongly correlated with the profiles of AZA (r = 0.57) and DEC (r = 0.68), suggesting that all three agents kill cancer cells primarily via DNMT1 degradation and DNA hypomethylation, rather than DNA damage from DNA-DNMT1 crosslinks (Fig. 1G, Extended Data Fig. 1J, and Supplementary Table 1).

Whereas cell killing by DNA-damaging agents tends to occur quickly, epigenetic killing mechanisms typically operate on longer timescales due to the gradual nature of chromatin remodeling. To test the hypothesis that these anti-cancer agents exhibit epigenetic drug kinetics, we modified the PRISM protocol to allow for a longer assay duration of 10 days and conducted PRISM timecourse experiments with >400 cell lines treated with GSK-3685032 or DEC. Consistent with an epigenetic killing mechanism, sensitivity to the GSK compound or DEC was time-dependent, with a significantly greater fraction of PRISM cell lines killed at later time points across both low and high doses of drug (Fig. 1H and Extended Data Fig. 1K). Validation in an AML cell line confirmed the time-dependent killing effects of DEC. In contrast, cancer cell killing by cytarabine, a structurally-related cytidine nucleoside analog that induces DNA damage without hypomethylation, did not exhibit such slow onset (Extended Data Fig. 1L).

### Discovery of genetic modifiers of DNMT1 inhibitors

To interrogate the specific pathways involved in tumor cell killing by DNA hypomethylation, we performed genome-wide CRISPR-Cas12a loss-of-function screens across three leukemia models treated with AZA or GSK-3685032 (Fig. 2A and Extended Data Fig. 2A-B). We hypothesized that if DNA hypomethylation, rather than DNA damage, is the primary killing mechanism, then drug sensitizers from the screen would primarily consist of epigenetic rather than DNA repair factors. The high quality of the screens was confirmed by the observation that killing by AZA was rescued by knockout of its bioactivating kinase, *Uridine Cytidine Kinase 2* (*UCK2*)^29^. In contrast, knockout of *UCK2* did not affect GSK-3685032 sensitivity, consistent with its phosphorylation-independent mechanism (Extended Data Fig. 2C-D). To ensure the generalizability of our screens, we analyzed shared hits across compounds, cell lines, and timepoints and identified 37 gene knockouts that sensitize cancer cells to DNMT1 inhibition (Fig. 2B-C and Supplementary Table 4). Indeed, these genes were predominantly epigenetic regulators with chromatin-binding functions, rather than DNA repair factors, and unbiased gene set enrichment analysis identified chromatin modification as the most enriched gene set (FDR < 2.87 × 10^−9^) (Supplementary Table 5). This further supports DNA hypomethylation as the primary mechanism of killing.

**Figure 2:**
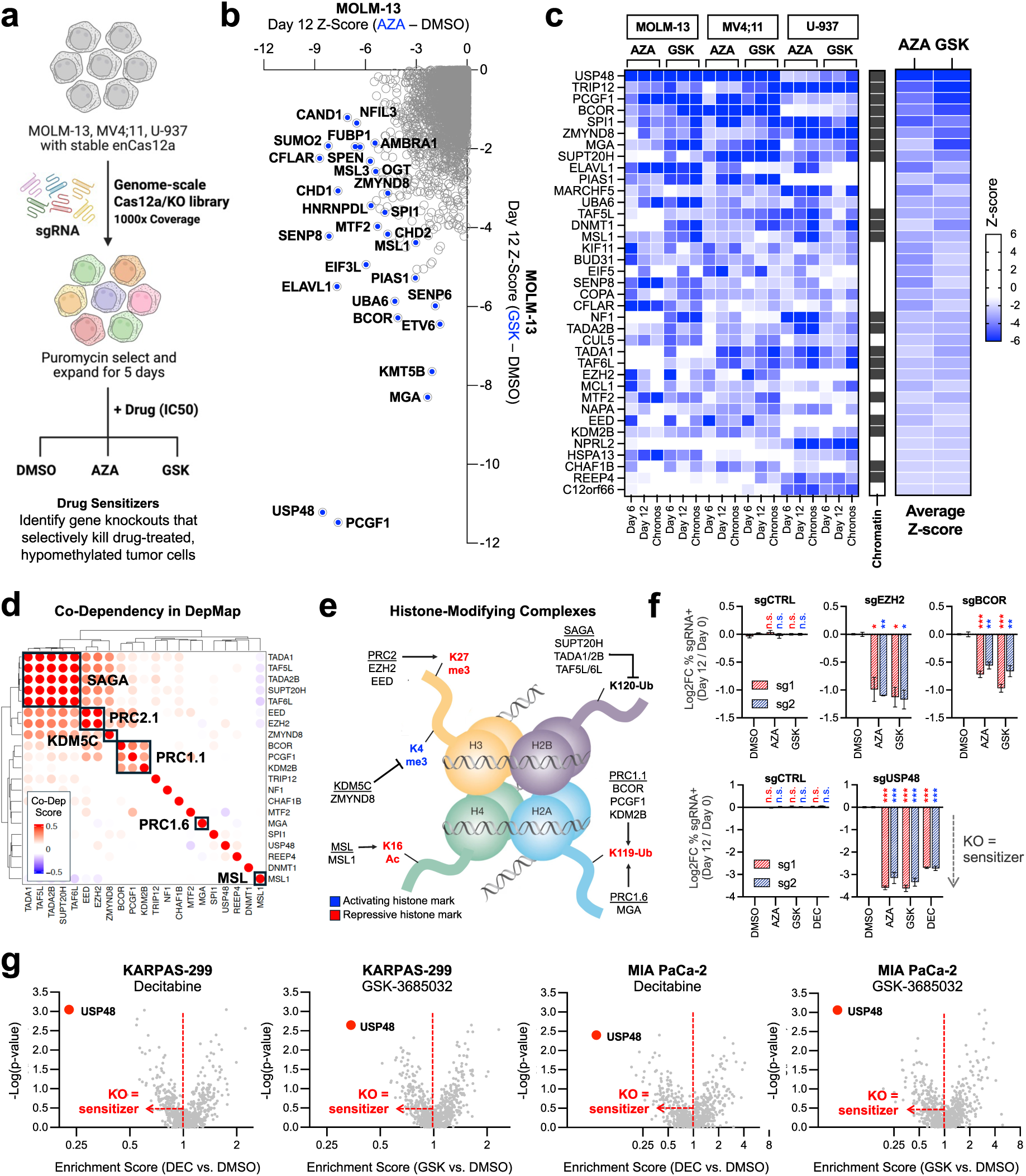
Chromatin-based modifiers of DNMT1 inhibitor anti-cancer activity. **a.** Genome-scale CRISPR-Cas12a screening strategy to identify genetic modifiers of DNMT1 inhibition. **b.** Z-score analysis of gene knockouts that sensitize MOLM-13 cells to AZA (n=2) compared to GSK-3685032 (n=2) at day 12. **c.** Heatmap of top drug sensitizers across different cell lines (MOLM-13, MV4;11, or U-937), timepoints (days 6 or 12), and anchor drugs (AZA or GSK-3685032). Chronos Z-scores represent a combined metric of gene knockout activity over time. Genes are ranked by average Z-score, and factors with known chromatin-based functions are annotated. **d.** Co-dependency analysis of top-scoring epigenetic hits from the CRISPR screens reveals distinct histone modifying complexes. A positive co-dependency score for a gene pair indicates that the individual gene knockouts produce a similar killing pattern across the >1,000 cell lines profiled in DepMap. **e.** Nucleosome schematic depicts histone modifications targeted by the chromatin complexes identified in the co-dependency analysis. **f.** MOLM-13 cell competition assays for validation of individual sgRNAs targeting components of PRC1.1 (*BCOR*) and PRC2.1 (*EZH2*) or a novel histone deubiquitinase (*USP48*). *EZH2* (n=2), *BCOR* (n=3), and *USP48* (n=3) knockout cells, but not cells expressing a control sgRNA (n=3), have a competitive disadvantage following treatment with a DNMT1 inhibitor compared to DMSO. **g.** CRISPR-Cas9 drug modifier screens with a ubiquitin-proteasome system focused sgRNA library in KARPAS-299 (lymphoid) or MIA PaCa-2 (pancreatic) cell lines treated with either DEC (n=3) or GSK-3685032 (n=3). ^∗^p < 0.05, ^∗∗^p < 0.01, ^∗∗∗^p < 0.001, determined by an unpaired, two-sided Student’s t-test. Mean values are shown unless otherwise specified, and error bars represent ± SEM.

### DNA hypomethylation creates dependency on histone-modifying enzymes

Strategies to enhance drug sensitivity require insight into the protective mechanisms by which tumor cells adapt to treatment. To elucidate how cancer cells adapt to DNA hypomethylation, we performed a co-dependency analysis across the >1,000 cell lines in the Cancer Dependency Map (DepMap). That is, we looked for pairwise correlation between epigenetic factors that we discovered as drug sensitizers to identify groups of genes with similar knockout killing profiles across the DepMap, indicating related biological function^33^. This analysis revealed five repressive chromatin complexes: PRC1.1 (BCOR, PCGF1, KDM2B), PRC1.6 (MGA), PRC2.1 (EED, EZH2), MSL (MSL1), and KDM5C (ZMYND8)^34,35^ (Fig. 2D). These complexes were differentially required in DNMT1 inhibitor-treated tumor cells, suggesting compensation for DNA hypomethylation through coordinated modulation of histone marks – either gaining repressive modifications (H3K27me3, H2K119Ub, or H4K16Ac) or losing activating marks (H3K4me3) (Fig. 2E). Genetic validation studies confirmed this model, as deletion of *BCOR* or *EZH2* sensitized leukemia cells to multiple DNMT1 inhibitors (Fig. 2F).

The histone DUB USP48^36,37^ emerged as a particularly striking and previously unrecognized chromatin-associated modifier of DNMT1 inhibitor activity. Multiple sgRNAs targeting *USP48* were selectively depleted following DNMT1 inhibitor treatment, but were largely maintained in DMSO-treated cells (Fig. 2F and Extended Data Fig. 3A). To more precisely define the role of USP48 in drug response, compared to other deubiquitinating enzymes, we conducted focused CRISPR modifier screens targeting 693 known components of the ubiquitin proteasome system, including DUBs and ubiquitin ligases^38^. *USP48* knockout again emerged as the top sensitizing mechanism, for both DEC and GSK-3685032, and enhanced drug-induced apoptotic cell death in validation studies (Fig. 2G, Extended Data Fig. 3B, and Supplementary Table 6). Follow-up experiments showed that USP48 loss-of-function was selective – the drug potentiation effect was correlated to DNMT1 inhibitor potency^26^, and *USP48* deletion did not enhance the activity of DNA-damaging agents such as doxorubicin or cytarabine (Fig. 3A and Extended Data Fig. 3C).

**Figure 3:**
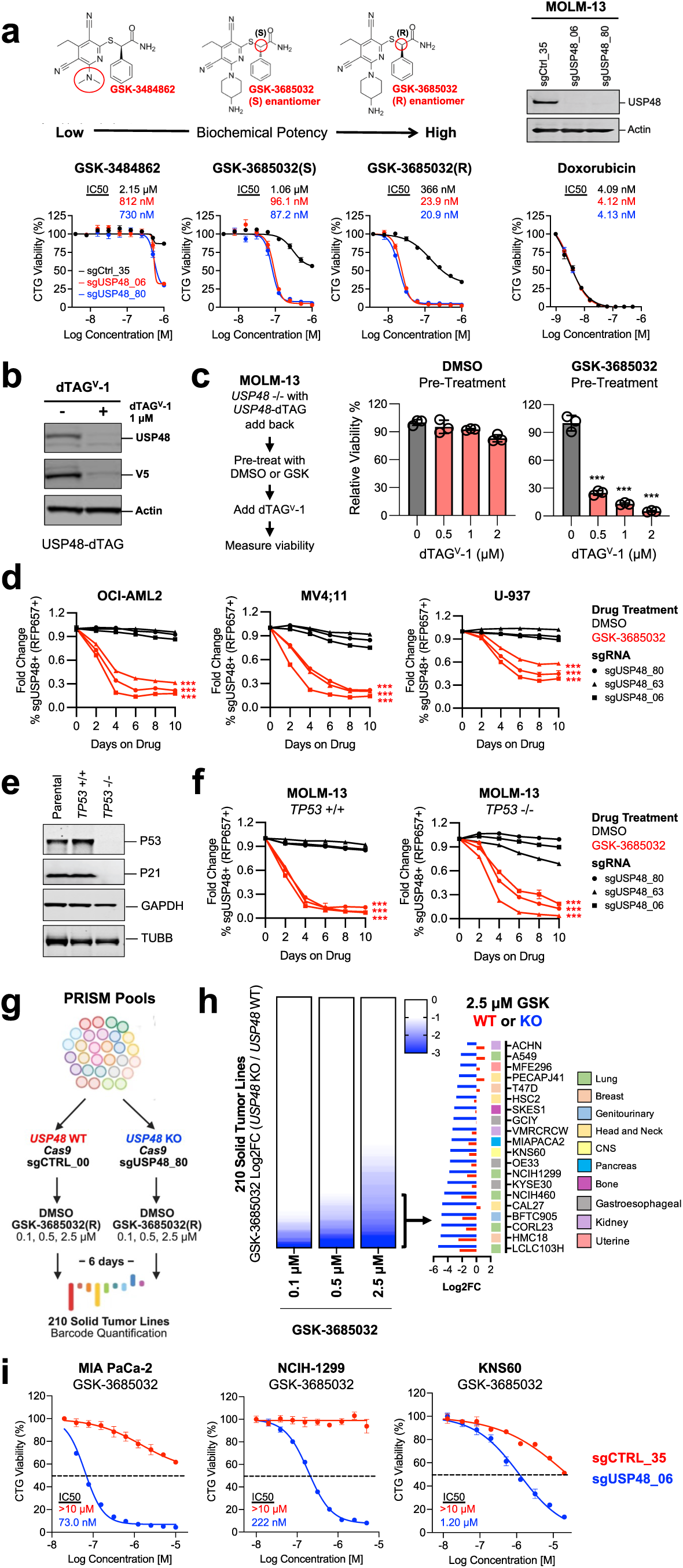
*USP48* deletion synergizes with DNMT1 inhibitors to induce tumor cell death. **a.** *USP48*-intact (sgCTRL_35) or *USP48*-deleted (sgUSP48_06 or sgRNA_80) MOLM-13 cells were validated for knockout by western blot and treated with compounds (n=3) in the GSK-3685032 chemical series for 72 hrs in a dose response. Compounds vary in their biochemical potency for DNMT1 inhibition. Doxorubicin (n=3) was used as a control for conventional DNA damage. **b.** Western blot validation of USP48 degradation in MOLM-13 cells with USP48-dTAG add-back. **c.** USP48-dTAG add-back cells were pre-treated with DMSO (n=3) or 1 μM GSK-3685032 (n=3) for 3 days to induce DNA hypomethylation, and dTAG^V^-1 (0, 0.5, 1, or 2 μM) was then added to induce USP48 degradation. Viable cells were counted following the addition of dTAG^V^-1. **d.** Competition assays with three unique pairs of control sgRNA (co-expression of *BFP*) and *USP48* sgRNA (co-expression of *RFP657*) in OCI-AML2 (*TP53*-intact), MV4;11 (*TP53*-intact), and U-937 (*TP53*-null) cells treated with DMSO (n=3) or 500 nM GSK-3685032 (n=3) over 10 days. A reduction in RFP657+ cells indicates a competitive disadvantage for *USP48* sgRNA-expressing cells compared to control sgRNA-expressing cells. **e.** Western blot analysis of P53 and P21 expression in isogenic *TP53* +/+ and *TP53* −/− MOLM-13 cells treated with 5 µM Nutlin-3 for 24 hrs. **f.** Competition assays with control sgRNA (co-expression of *BFP*) and *USP48* sgRNA (co-expression of *RFP657*) in isogenic *TP53* +/+ and *TP53* −/− MOLM-13 cells treated with DMSO (n=3) or GSK-3685032 (n=3). **g.** Experimental approach to test the combination effect of *USP48* deletion and GSK-3685032 (n=3) across 219 solid tumor cell lines in PRISM. **h.** Identification of a subset of solid tumor cell lines with greater sensitivity to the *USP48* knockout and GSK-3685032 combination compared to the single perturbations. **i.** Dose-response curves for isogenic *USP48*-intact (sgCTRL_35) or *USP48*-deleted (sgUSP48_06) MIA PaCa-2, NCIH-1299, or KNS60 cells after 6-day treatment with GSK-3685032 (n=3). ^∗^p < 0.05, ^∗∗^p < 0.01, ^∗∗∗^p < 0.001, determined by an unpaired, two-sided Student’s t-test. Mean values are shown unless otherwise specified, and error bars represent ± SEM.

To better approximate the effect of USP48 inhibition that might be induced by a drug, we engineered a C-terminal dTAG degron^39,40^ into USP48, thereby allowing for its conditional degradation after treatment with the dTAG^V^-1 compound (Fig. 3B). As with complete genetic knockout, USP48 protein degradation led to AML cell death only in the presence of a DNMT1 inhibitor (Fig. 3C). Furthermore, the effect of USP48 loss-of-function as a drug sensitizer was consistent across multiple models of AML, independent of *TP53* mutation status^41^ (Fig. 3D-F and Extended Data Fig. 3D).

To test the broad applicability of *USP48* deletion to potentiate DNMT1 inhibitor activity in solid tumors, we transduced 219 cell line models in PRISM with a *USP48* sgRNA or a control sgRNA, together with co-expression of *Cas9*, resulting in *USP48*-knockout or *USP48*-intact cell line pools (Fig. 3G). *USP48* deletion markedly sensitized a subset of solid tumor lines to DNMT1 inhibition, dramatically reducing the IC50 for GSK-3685032 to nanomolar concentrations (Fig. 3H-I and Supplementary Table 7). Thus, *USP48* deletion sensitizes cancer cells across diverse lineages to death from drug-induced DNA hypomethylation.

### Nuclear localization and catalytic activity are required for USP48 function

While many DUBs function through catalytic cleavage of ubiquitin marks on substrate proteins, others have lost catalytic activity over the course of evolution and function mainly through protein-protein interactions^42^. These properties have not been reported for USP48. Amino acid conservation analysis showed that the USP48 catalytic domain is conserved from yeast to humans (Extended Data Fig. 4A). In addition, the AlphaFold2 (AF2) structural model^43^ of USP48 revealed an intact deubiquitinase domain with homology to the catalytically-active DUB USP7 (root-mean-square deviation=3.8Å), including a properly oriented catalytic triad (Cys-98, Asn-370, His-353)^44^ (Fig. 4A-B). *In vitro* DUB assays demonstrated that USP48 efficiently cleaved K63-linked tetra-ubiquitin chains, a process blocked by the 17e^45^ and PR-619^46^ DUB inhibitors, further indicating that USP48 has deubiquitinating enzymatic function^47^ (Fig. 4C).

**Figure 4:**
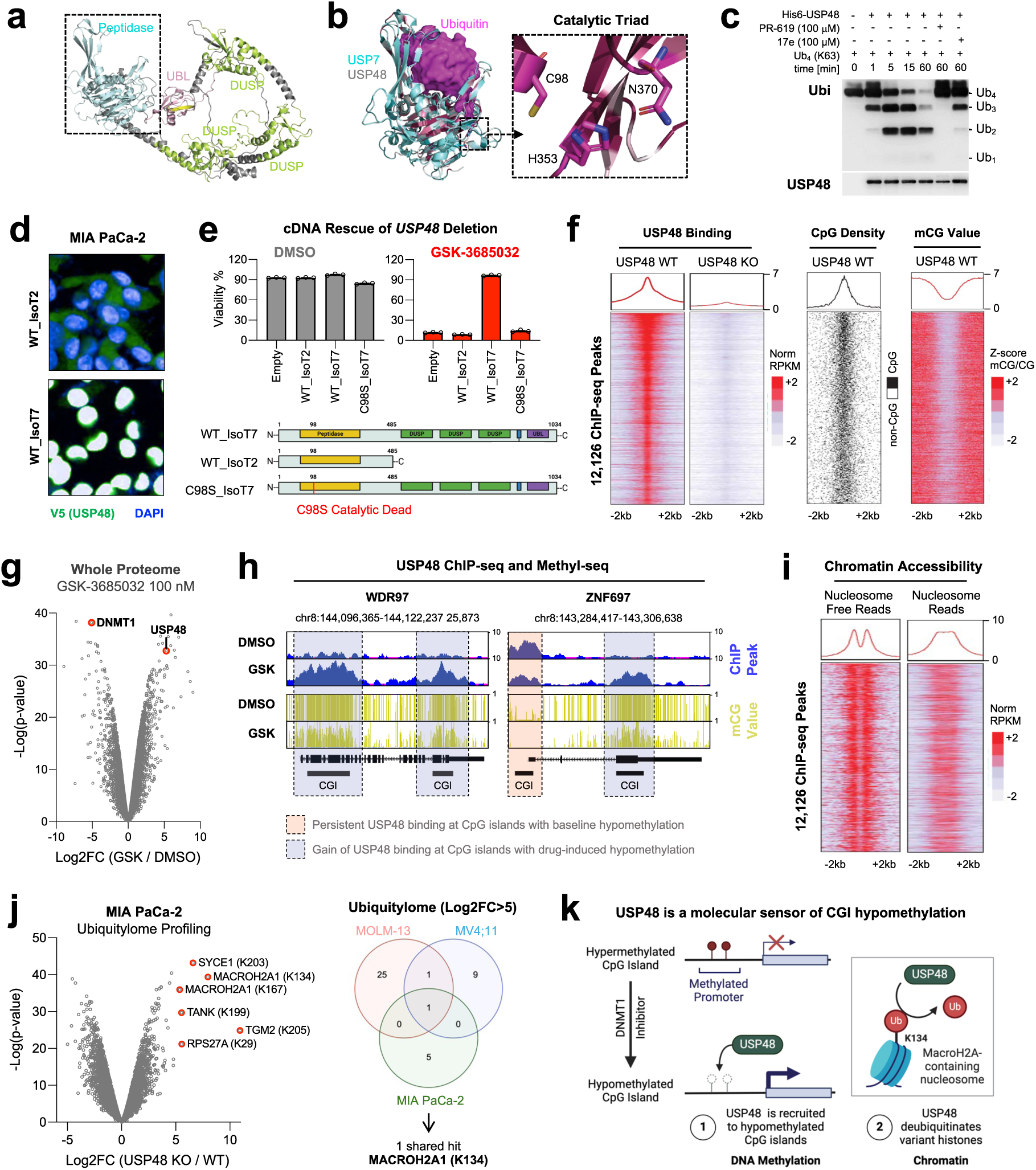
USP48 is a molecular sensor linking DNA hypomethylation and histone deubiquitination. **a.** AF2 model of full-length human USP48 protein. The N-terminal peptidase domain, three DUSP domains, and a C-terminal ubiquitin-like (UBL) domain are highlighted in cyan, green, and pink, respectively. **b.** Superposition of the catalytic domains of USP48 (rainbow) and USP7 (cyan) in complex with ubiquitin (magenta surface) (PDB: 5KYE). The well-conserved catalytic triad (Cys-98, His-353, and Asn-370) is illustrated. **c.** *In vitro* K63-linked tetra-ubiquitin kinetic DUB assay with His6-USP48. Addition of DUB inhibitors (PR-619 or 17e) blocks tetra-ubiquitin hydrolysis. **d.** Immunofluorescence images depicting cytoplasmic localization of the short USP48 isoform (WT_IsoT2) and nuclear localization of long USP48 isoform (WT_IsoT7) in MIA PaCa-2 cells. **e.** Rescue of *USP48* deletion with *WT_IsoT7* in MIA PaCa-2 cells after treatment with GSK-3685032 (n=3). *WT_IsoT2* and a catalytic dead version of the long *USP48* isoform (*C98S_IsoT7*) fail to rescue. **f.** Heatmaps of USP48 ChIP-seq peaks in *USP48*-intact (sgCTRL_35) (n=2) and *USP48*-deleted (sgUSP48_80) (n=2) MOLM-13 cells. Additional heatmaps show CpG density and mCG content across 4-kb windows centered on the identified USP48 ChIP-seq peaks. USP48 selectively binds hypomethylated CpG islands. **g.** Whole proteome profiling of MOLM-13 cells treated with 100 nM GSK-3685032 (n=2) compared to DMSO (n=2). **h.** Genome browser tracks for USP48 ChIP-seq peaks (blue) and mCG values (yellow) in MOLM-13 cells treated with DMSO or 1 μM GSK-3685032. USP48 localizes to newly hypomethylated CpG islands in GSK-3685032-treated cells. **i.** ATAC-seq profiles in MOLM-13 cells (n=2) showing predicted nucleosome enrichment across 4-kb windows for the identified USP48 binding sites. **j.** Ubiquitylome (normalized to proteome) profiling in MIA PaCa-2 cells reveals increased ubiquitination of the variant histone MACROH2A1 in *USP48*-deleted (sgUSP48_06) (n=3) cells compared to *USP48*-intact (sgCTRL_35) (n=3) cells. Venn diagram illustrates ubiquitinated proteins (and target lysine sites) that are upregulated (Log2FC>5) in the ubiquitylome following *USP48* deletion (sgUSP48_06), and the intersection across three cell lines (MIA PaCa-2, MV4;11, and MOLM-13). MACROH2A1 Lys-134 is the single shared target across the three cell models. **k.** Mechanistic model: USP48 functions as a molecular sensor of DNA hypomethylation. USP48 is specifically recruited to hypomethylated CpG islands where it deubiquitinates non-canonical histones, most notably MACROH2A1, thereby providing a molecular link between DNA hypomethylation and histone modification.

To determine whether catalytic activity is required for USP48’s role in modulating response to DNA hypomethylation, we performed *USP48* mutational analysis, asking whether *USP48* mutant constructs could rescue cells from GSK-3685032 treatment. These experiments showed that nuclear localization and an intact catalytic domain of USP48 are essential, and moreover, a catalytically-dead C98S mutant failed to rescue cells from GSK-3685032-induced cell death (Fig. 4D-E).

### USP48 operates as a CpG island methylation sensor

To investigate the chromatin-based function of USP48, we performed chromatin immunoprecipitation sequencing (ChIP-seq). We identified 12,126 high-confidence USP48 binding sites that were lost in *USP48*-knockout cells and observed that USP48 occupancy was restricted to hypomethylated CpG islands (CGIs) (Fig. 4F, Extended Data Fig. 4B, and Supplementary Table 8). USP48 protein levels increased rapidly following GSK-3685032 treatment with concomitant binding to newly hypomethylated CGI sites, consistent with a protective feedback mechanism (Fig. 4G-H, Extended Data Fig. 4C, and Supplementary Table 3). Additionally, integration with an assay for transposase accessible chromatin sequencing (ATAC-seq) showed that USP48 preferentially associated with nucleosome-occupied regions, suggesting that its localization is mediated by nucleosome binding (Fig 4I, Extended Data Fig. 4D, and Supplementary Table 9).

We confirmed that DNMT1 itself is not a substrate of USP48 in that DNMT1 stability remained unaltered in *USP48*-deleted cells (Extended Data Fig. 4E-F and Supplementary Table 3). Proteome-wide ubiquitylome profiling, however, revealed that *USP48* deletion led to increased ubiquitination of specific lysine residues on several non-canonical histone proteins – particularly MACROH2A1^48^ (Lys-134) – representing putative USP48 substrates (Fig. 4J, Extended Data Fig. 4G, and Supplementary Table 3). Direct interaction between USP48 and MACROH2A1 was confirmed by co-immunoprecipitation followed by mass spectrometry (Extended Data Fig. 4H-I and Supplementary Table 10).

Collectively, these findings establish USP48 as a molecular sensor of DNA hypomethylation: USP48 is induced and selectively recruited to nucleosomes at hypomethylated CGIs, where it deubiquitinates non-canonical histones. This previously unrecognized chromatin-based function of USP48 establishes the mechanistic link between DNA methylation and histone modification, and thereby explains the synergy between USP48 loss-of-function and DNMT1 inhibitor activity (Fig. 4K).

### *USP48* copy number loss is frequent in human tumors and sensitizes to DNMT1 inhibition *in vivo*

Having established that *USP48* knockout sensitizes leukemia and solid tumor cell lines to treatment with DNMT1 inhibitors, we next asked whether any human tumors have naturally-occurring *USP48* gene deletions that might similarly create synthetic lethal treatment opportunities. Using publicly available datasets representing 105,260 patients^49^, we found evidence of heterozygous or homozygous *USP48* deletions across a wide range of tumor types with a deletion frequency of 15% overall (Fig. 5A and Copy Number Alteration Analysis in Methods). *USP48* copy number loss was in part driven by larger deletions at the tumor suppressive 1p36 locus, which is recurrently lost in a variety of cancer types. Indeed, previous reports indicate that biallelic deletions of *USP48* occur in up to 18% of meningiomas^50^, 11% of pancreatic adenocarcinomas^51^, 5% of colon adenocarcinomas^52^, 3% of breast cancers^53^, 1.8% of metastatic melanomas^54^, and 1.3% of pediatric AML^55^. This result suggests that such tumors might be particularly sensitive to DNMT1 inhibition.

**Figure 5:**
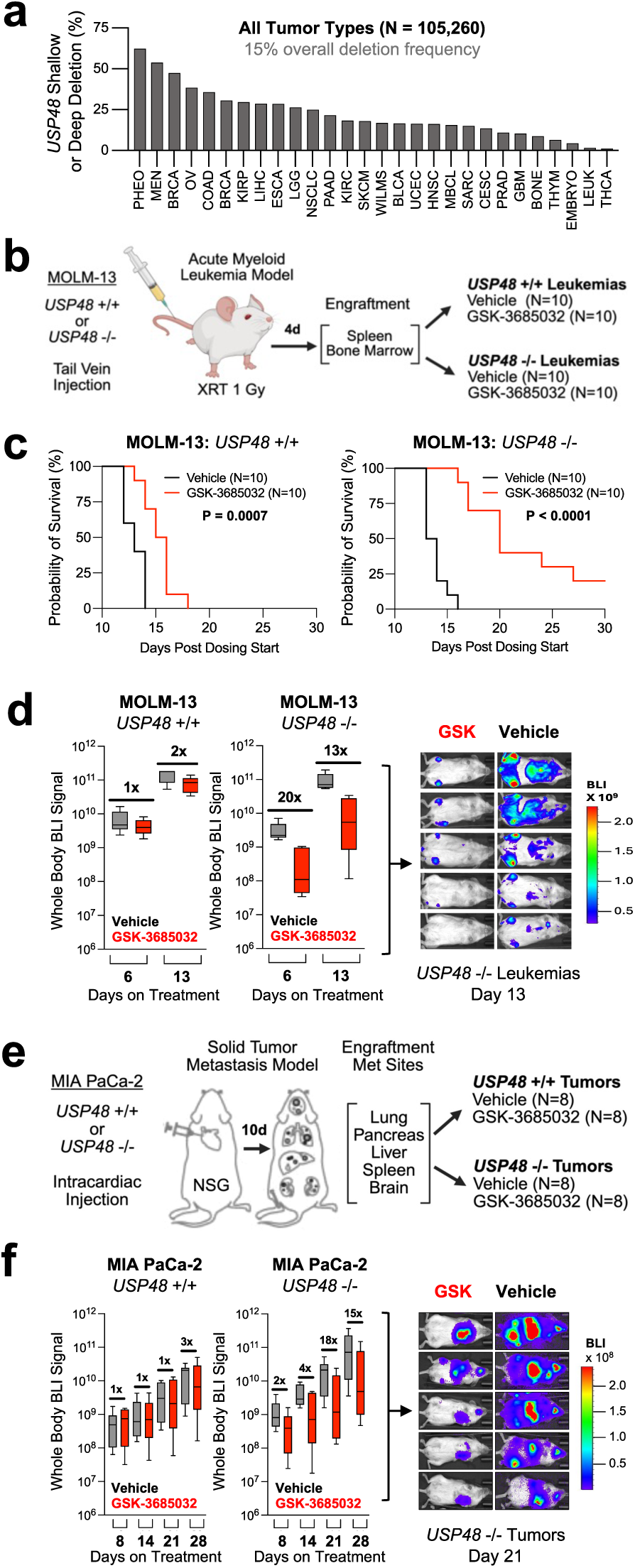
*USP48* copy number loss is frequent in human tumors and sensitizes to DNMT1 inhibition *in vivo*. **a.** Frequency of shallow or deep deletions of *USP48* across non-redundant human tumor genomic datasets in cBioPortal. **b.** MOLM-13 systemic AML model of *USP48* +/+ or *USP48* −/− leukemias treated with vehicle or GSK-3685032 (30 mg/kg BID). **c.** Kaplan-Meier survival analysis of *USP48* +/+ or *USP48* −/− MOLM-13 leukemias treated with vehicle or GSK-3685032 (30 mg/kg BID). **d.** Left panel: BLI measurements of all mice in the *USP48* +/+ or *USP48* −/− MOLM-13 leukemia cohorts while on treatment. Right panel: BLI images for 5 representative mice in the *USP48* −/− cohort at day 10 after treatment initiation. **e.** MIA PaCa-2 pancreatic cancer metastasis model. **f.** Left panel: BLI measurements for all mice in the *USP48* +/+ or *USP48* −/− MIA PaCa-2 tumor cohorts while on treatment with vehicle or GSK-3685032. Right panel: BLI images for 5 representative mice in the *USP48* −/− cohort at day 21 after treatment initiation.

To test the hypothesis that *USP48* loss-of-function alterations might sensitize human tumors to DNMT1 inhibitor treatment *in vivo*, we conducted tumor xenograft experiments using human cancer cells grown in immunodeficient mice. We first explored an AML model^56^, wherein luciferase-expressing MOLM-13 cells (*USP48* knockout or wildtype) were transplanted into sublethally-irradiated NOD SCID gamma (NSG) mice via tail vein injection, enabling engraftment in both the bone marrow and spleen (Fig. 5B). *USP48*-deleted leukemias demonstrated marked sensitivity to GSK-3685032, with a 20-fold reduction in bioluminescence imaging (BLI) signal and prolonged overall survival (Fig. 5C-D). In contrast, *USP48*-intact leukemias showed only a modest response.

Given the preponderance of *USP48* biallelic deletions in pancreatic cancer, we next established a metastatic model using luciferase-expressing MIA PaCa-2 pancreatic cells administered via intracardiac injection to ensure widespread dissemination^57^ (Fig. 5E). BLI confirmed extensive metastatic tumor formation in the pancreas, liver, lungs, and brain, consistent with the pattern of metastasis seen in patients with advanced pancreatic cancer (Extended Data Fig. 5A). Mice bearing *USP48*-deleted metastatic tumors showed an 18-fold reduction in disease burden and increased survival after three weeks of GSK-3685032 treatment, whereas *USP48*-intact tumors were resistant to the drug *in vivo* (Fig. 5F and Extended Data Fig. 5B). These results demonstrate that biallelic *USP48* deletion sensitizes leukemias and metastatic solid tumors to DNMT1 inhibition *in vivo*. This supports *USP48* copy number status as a potential predictive biomarker for clinical response to DNMT1 inhibitors.

## DISCUSSION

The work described here addresses a long-standing question in clinical oncology: how do the commonly used drugs AZA and DEC kill leukemia cells, and are there opportunities to extend their use to solid tumors? Our findings reveal new insights into the mechanisms of tumor cell killing by these drugs, clearly pointing to their primary function as DNA hypomethylating agents rather than as DNA-damaging agents, a mechanism originally intended for these drugs. Using functional genomic approaches and new chemical tools, including the non-DNA-damaging DNMT1 inhibitor GSK-3685032, we demonstrate that DNA hypomethylation and an adaptive epigenetic response are key determinants of the anti-tumor activity of these agents. These data also address an important historical observation from the clinic, namely the prolonged time to clinical responses seen in leukemia patients treated with low dose AZA or DEC^58^. Our data reveal a mechanistic understanding of the epigenetic remodeling that occurs under such treatment regimens.

This study also provides a window into how tumor cells sense and adapt to abrupt loss of DNA methylation. Through a series of CRISPR drug modifier screens across a diversity of cancer models, we discovered a network of epigenetic factors that becomes essential in hypomethylated cancer cells. This network comprises canonical repressive histone modifying complexes, including the polycomb complexes PRC1.1 and PRC2.1, that serve as compensatory mechanisms when DNA methylation is lost. Our data are consistent with the recent report of synergistic activity between DNMT1 inhibitors and PRC2 loss-of-function mutations^59^. Overall, we discovered intimate coordination between DNA methylation and histone modifications in maintaining epigenetic homeostasis. Central to these findings is the discovery of USP48 as a molecular sensor of DNA hypomethylation – a previously unknown mechanism in which USP48 is rapidly recruited to hypomethylated CpG islands, where it deubiquitinates non-canonical histones, directly coupling DNA hypomethylation and histone ubiquitination. DNA methylation and histone modification have long been known to be the two principal mechanisms by which gene expression is epigenetically regulated. However, it has been unknown how this dynamic system establishes homeostatic balance. USP48 appears to be central to this balance, allowing for dynamic adaptive responses to changes in DNA methylation state.

These mechanistic insights have important therapeutic implications. First, they challenge the conventional view that DNMT1 inhibitor activity is limited to hematological malignancies, suggesting their broader utility across many cancer types, especially when guided by predictive biomarkers. Specifically, our discovery of *USP48* as a synthetic lethal dependency in hypomethylated cancer cells provides a potential patient selection strategy: human tumors with naturally occurring *USP48* deletions are predicted to have enhanced sensitivity to DNMT1 inhibition. Homozygous deletions of *USP48* are observed in several types of human solid tumors, and could easily be detected in the clinical setting either by DNA sequencing or fluorescence in situ hybridization (FISH). In addition, our finding of multiple chromatin-based dependencies in hypomethylated tumor cells suggests rational combination strategies targeting both DNA methylation and histone modifications. Such an approach will require highly selective epigenetic modulators, including further optimization of the selectivity and potency of existing DUB inhibitors^60^.

Collectively, our work establishes DNA hypomethylation through DNMT1 degradation as the primary anti-cancer killing mechanism for the clinically approved drugs AZA and DEC, and uncovers a previously unrecognized epigenetic regulatory circuit centered on histone-modifying enzymes such as USP48. The work further suggests that DNMT1 inhibitors should be explored in clinical trials in patients with solid tumors harboring *USP48* deletions.

## METHODS

### Cell Culture and Lentiviral Transduction

Cell culture conditions and antibiotic concentrations for each cell line are described in detail in the Supplementary Information. A modified spinfection protocol was used for lentiviral transduction of cancer cell lines for stable expression of individual sgRNAs and cDNAs. Briefly, cells resuspended in media and polybrene (10 μg/mL) were mixed with an equal volume of lentivirus-containing medium. Immediately following spinfection (2000 RPM, 90 min, 37°C), fresh media was added. The cells were incubated overnight and then split accordingly. Transductions of PRISM cell line pools were performed using the same spinfection protocol.

### Drug Treatments and Dose-Response Analysis

For dose-response viability assessment, individual cell lines were resuspended in media and plated (200 µl/well) in black, clear bottom 96-well plates (Corning, 3904) at a cell density optimized for log-phase growth for the duration of the experiment, which was typically 3 or 6 days. For suspension cell lines, compound was added the same day as plating. For adherent cell lines, compound was added the day after cell plating. The Cell Titer-Glo Luminescent Cell Viability Assay (G7571, Promega) was used for viability measurements, according to the manufacturer’s protocol. Luminescence values for each drug dose were normalized to the value of the corresponding DMSO-only control, and the average normalized luminescence and SEM are plotted for each dose (n=3). Dose-response curves were fitted to log(inhibitor) vs. response (variable slope, four parameters) in GraphPad PRISM with reported IC50 values based on the best-fit values. A full list of the compounds and compound vendors used in the study are listed in the Supplementary Information.

### PRISM Viability Screening

For the 6-day PRISM assay, 766 DNA-barcoded human cancer cell lines were distributed across 7 uber pools (46-144 lines per uber pool) and expanded prior to screening^27^. The PRISM uber pools were then counted on the Vi-CELL BLU Cell Viability Analyzer (Beckman-Coulter) and plated on 24-well poly-D-lysine-coated plates (Corning, 356414) in RPMI-1640 without phenol red (Gibco, 11835030) supplemented with 10% fetal bovine serum (FBS) (Sigma-Aldrich, F7524) and 1% penicillin/streptomycin (P/S) (Gibco, 15140122). Plating density for each PRISM uber pool was optimized based on estimated doubling time. After 24 hrs, compounds were added in an 7-point dose response format (n=3) using the D300e Digital Dispenser (Tecan, 30100152) and refreshed every 48 hrs. Compound doses were as follows: AZA (0.25, 0.5, 1, 2.5, 5, 10, and 20 µM), DEC (0.05, 0.1, 0.25, 0.5, 1, 2.5, and 5 µM), and GSK-3685032 (0.25, 0.5, 1, 2.5, 5, 10, and 20 µM) in addition to a DMSO-only control. The proteosome inhibitor bortezomib and the BRAF inhibitor vemurafenib served as positive control compounds. Following 6 days of drug treatment, media was gently aspirated and cells were lysed directly in plates using buffer containing 20 mM Tris-HCl (pH 8.4), 50 mM KCl, 0.45% Tween-20 (Sigma, P9416), 0.45% NP40 (Sigma, I8896), 10% Proteinase K (Qiagen, 19133), and nuclease-free water (Invitrogen, 10977015). Plates were covered with foil and heated for 60 min at 60°C followed by long-term storage at −80°C. Protocol modifications for the extended timecourse and CRISPR/Cas9 PRISM screens, library preparation, next generation sequencing, and data analysis methods are described in the Supplementary Information.

### Humagne CRISPR-Cas12a Drug Modifier Screens

The Humagne genome-scale CRISPR-Cas12a sgRNA library was obtained from the Broad Institute Genetic Perturbation Platform (GPP). This library consists of Sets A and B, each of which targets the same 19,755 genes. Each set has 1 vector per gene, and each vector contains 2 unique sgRNAs targeting the same gene^61^. Lentivirus for Humagne Sets A and B was obtained from GPP and titrated to a goal infection efficiency of 0.3–0.5. MOLM-13, MV4;11, or U-937 cells with stable expression of enhanced *Cas12a* derived from *Acidaminococcus sp*. (*enAsCas12a*) were generated via lentiviral transduction of the parental cell lines with pRDA_174 (Addgene, 136476) followed by balastocidin selection (8-10 µg/mL). For each cell line, >70% Cas12a cutting activity was confirmed using the pRDA_221 Cas12a activity assay (Addgene, 169142). On the first day of the screen, MOLM-13-Cas12a, MV4;11-Cas12a, or U-937-Cas12a cells were infected with Humagne Sets A or B virus in 12-well plates via centrifugation at 2000 RPM at 37°C to achieve 1000x coverage of each library set. The following day, the cells were split into 2 replicate flasks for Set A and 2 replicate flasks for Set B, selected with 2-4 µg/mL puromycin, and then expanded for 5 days. Replicates were then seeded into flasks containing DMSO, AZA, or GSK-3685032. The following compound doses (in addition to a DMSO-only control) were used for each screen: MOLM-13 (1 µM AZA and 0.5 µM GSK-3685032), MV4;11 (1.5 µM AZA and 0.5 µM GSK-3685032), and U-937 (1 µM AZA and 0.5 µM GSK-3685032). For the duration of the screen, cells were maintained at 37°C and 5% CO2 in T-175 flasks (Corning) in RPMI with 20% FBS and 1% P/S. Cells were reseeded with fresh media and drug every 3 days with a minimum of 22E6 cells replated for each replicate per passage (to maintain ∼1000x library representation). Cell pellets were collected before drug was added (input sample) and after 6 or 12 days of drug exposure (day 6 or 12 samples). Genomic DNA was isolated from cell pellets using the NucleoSpin Blood XL Columns (Macherey Nagel, 740950.50S). Genomic DNA PCR and sequencing was performed by GPP, and the data was analyzed as described in the Supplementary Information.

### USP48 Structural Modeling

Structural models of full-length USP48, the peptidase domain, the catalytic triad, and ubiquitin binding were generated with AF2. To assess for structural homology between USP48 and the related DUB USP7, we superposed the catalytic domains of USP7 (PDB: 5KYE, residues 209-553) and USP48 (AF2 USP48 model, residues 89-421) using Molecular Operating Environment (MOE) 2020.0901 from Chemical Computing Group and performed a root-mean-square deviation (RMSD) analysis. For evolutionary conservation assessment, we used the ConSurf public portal (https://consurf.tau.ac.il/consurf_index.php)^62^.

### Methylation-Sequencing

MOLM-13 cells were plated at 200,000 cells/mL in 6-well dishes and treated with DMSO (n=1) or 1 μM GSK-3685032 (n=1) for 72 hrs with daily refreshment of drug. Cells were then pelleted by centrifugation and washed with PBS followed by genomic DNA extraction using the QIAamp Blood & Cell Culture DNA Mini Kit (Qiagen, 51304). DNA was quantified using the Qubit dsDNA BR Quantitation Kit (Thermo Fisher Scientific, Q32850) and submitted to Azenta (South Plainfield, NJ, USA) for Methyl-seq enzymatic treatment, library preparation, and methylation sequencing, as described in the Supplementary Information.

### ChIP-Sequencing

*USP48* wildtype and *USP48* knockout MOLM-13 lines were generated by lentiviral transduction of MOLM-13-Cas9 cells with pLentiGuide-Puro (Addgene, 52963) containing either sgCTRL_35 (n=2) or sgUSP48_80 (n=2), followed by puromycin (4 μg/mL) selection. For the drug treatment conditions, parental MOLM-13 cells were treated with either DMSO (n=2) or 1 μM GSK-3685032 (n=2) for 48 hrs. All cross-linking, nuclear isolation, immunoprecipitation, library preparation, and sequencing steps were performed as described previously^63^ and in the Supplementary Information.

### ATAC-Sequencing

MOLM-13 cells (n=2) were plated at 100,000 cells/mL in 6-well dishes and approximately 4 days later, cells were pelleted by centrifugation, resuspended in Recovery Cell Culture Freezing Medium (Gibco, 12648010), and frozen in cryotubes (Gibco). Tubes of viably frozen cells on dry ice were submitted to Azenta (South Plainfield, NJ, USA) for Tn5 treatment, library preparation, and sequencing, as outlined in the Supplementary Information.

### Global Proteome and Ubiquitylome Profiling

The *USP48* wildtype and *USP48* knockout MOLM-13 (n=2), MV4;11 (n=3), and MIA PaCa-2 (n=3) lines were generated by transducing *Cas9*-expressing cells with pLentiGuide-Puro (Addgene, 52963) containing either sgCTRL_35 or sgUSP48_06, followed by puromycin (1-4 µg/mL) selection. All drug treatments were for approximately 48 hrs, and the following compounds and doses were used: MOLM-13 (0.1, 0.25, and 0.75 µM GSK-3685032) (n=2), MV4;11 (0.25 µM GSK-3685032) (n=3), and MIA PaCa-2 (1.5 µM GSK-3685032) (n=3). Cells were pelleted by centrifugation, pellets were washed 3 times in ice-cold PBS, and samples were then flash frozen on dry ice plus ethanol and stored at −80°C prior to processing. Lysis and digestion, enrichment of K-ε-GG peptides, TMT labeling, LC-MS, and data analysis were performed as described in the Supplementary Information.

### *USP48* Copy Number Analysis

Using the cBioPortal for Cancer Genomics (https://www.cbioportal.org/), a curated set of non-redundant cancer genomic studies was analyzed for *USP48* copy number alterations, yielding a dataset with 105,260 patients across a diversity of tumor types. Deep and shallow deletions were queried using the terms HOMDEL or HETLOSS, respectively. The percentage of patients with predicted deep or shallow *USP48* deletions was calculated for each tumor type, and studies with fewer than 100 patients were excluded from the analysis. Individual genomic sequencing studies were then queried to establish the estimated percentage of biallelic deletions for select tumor types.

### *In Vivo* Murine Tumor Xenograft Studies

For the systemic AML model, NSG mice (Jackson Laboratories) were irradiated (1 Gy) and then transplanted 24 hrs later with 100,000 Firefly luciferase-expressing *USP48* wildtype (sgCTRL_20) or *USP48* knockout (sgUSP48_80) MOLM-13 cells via IV tail vein injection. GSK-3685032 (30 mg/kg SQ BID, resuspended in 10% captisol adjusted to pH 4.5-5 with 1M acetic acid) or vehicle treatment was initiated 4 days after cell injection and continued for a total of 30 days (n=10 for each group). Endpoints of the study included total body BLI signal and overall survival. For the pancreatic cancer metastasis model, 2E6 Firefly luciferase-expressing *USP48* wildtype (sgCTRL_20) or *USP48* knockout (sgUSP48_80) MIA PaCa-2 cells were transplanted via ultrasound-guided intracardiac injection, as described previously^57^. Dosing with GSK-3685032 (30 mg/kg SQ BID, resuspended in 10% captisol adjusted to pH 4.5-5 with 1M acetic acid) or vehicle was initiated 10 days after cell injection to ensure adequate metastatic tumor formation, and treatment was continued for a total of 14 days (n=8 for each group). Total body BLI signal and overall survival were monitored. The logrank (Mantel-Cox) test was used to assess for statistically-significant differences in survival between the vehicle-treated and GSK-3685032-treated cohorts. All *in vivo* murine studies were conducted under the IACUC animal protocol 08-023 at the Lurie Family Imaging Center at the Dana-Farber Cancer Institute.

## Data Availability

Raw data (FASTQ files) and processed data (WIG files) for the epigenomic profiling datasets (Methyl-seq, ChIP-seq, and ATAC-seq) generated in this study are available on Gene Expression Omnibus (GEO) under accession number GSE295429. The following secure token has been created to allow reviewers to access the raw GEO data while it remains private: ulqpoqugtrwrzc. For the proteomics studies (global proteome, ubiquitylome, and IP-MS), the original mass spectra and the protein sequence databases used for searches have been deposited in the public proteomics repository MassIVE (http://massive.ucsd.edu) and are accessible at ftp://MSV000097907@massive-ftp.ucsd.edu when providing the dataset password: DNMT1. If requested, provide the username: MSV000097907. These datasets will be made public upon acceptance of the manuscript.

## Supplementary Methods

A detailed description of additional methods and reagents used in this study, including lentiviral expression vectors, sgRNA sequences, and cDNA sequences, is presented in the Supplementary Information.

## ACKNOWLEDGEMENTS

We thank Ben Ebert, Rich Stone, Brad Bernstein, Brian Liau, Bruce Chabner, Tim Graubert, Joanna Morris, Ruth Densham, Kevin Ngan, and Steven Corsello for helpful scientific discussions. All *in vivo* murine experiments were performed by the Lurie Family Imaging Center at the Dana-Farber Cancer Institute by Q.D. Nguyen, N. Kania, and Z. Sall. This work was funded by the National Cancer Institute (NCI) grant 1R35CA242457–01 (T.R.G.), NCI Paul Calabresi Award for Clinical Oncology 5K12CA087723-22 (R.V.P.), Edward P. Evans Foundation Young Investigator Award (R.V.P.), and the National Institutes of Health T32 institutional training grant 2T32HL116324-06 (R.V.P.). The proteomic studies were supported in part by grants U24CA270823 and U01CA271402 (S.A.C.) from the NCI Clinical Proteomic Tumor Analysis Consortium program, as well as a grant from the Dr. Miriam and Sheldon G. Adelson Medical Research Foundation (N.D.U. and S.A.C.).

## AUTHOR CONTRIBUTIONS

R.V.P. and T.R.G conceived of the project and funded the work. R.V.P., T.R.G, J.W., N.U., S.A.C, J.A.R., J.G.D., M.S., M.K., and M.N. designed and supervised the research. R.V.P, Q.Y., Y.L., J.C.R, D.B., M.C.S., L.M., M.D., K.N., D.L.B., H.B.W., C.T., A.G., M.Y.C., Q.G., D.R.M., M.M.R, M.G.R., and B.I. performed the research. Q.Y. and J.W. led the epigenomic analyses. R.V.P. and T.R.G wrote the manuscript.

## EXTENDED DATA FIGURES

**Extended Data Figure 1:**
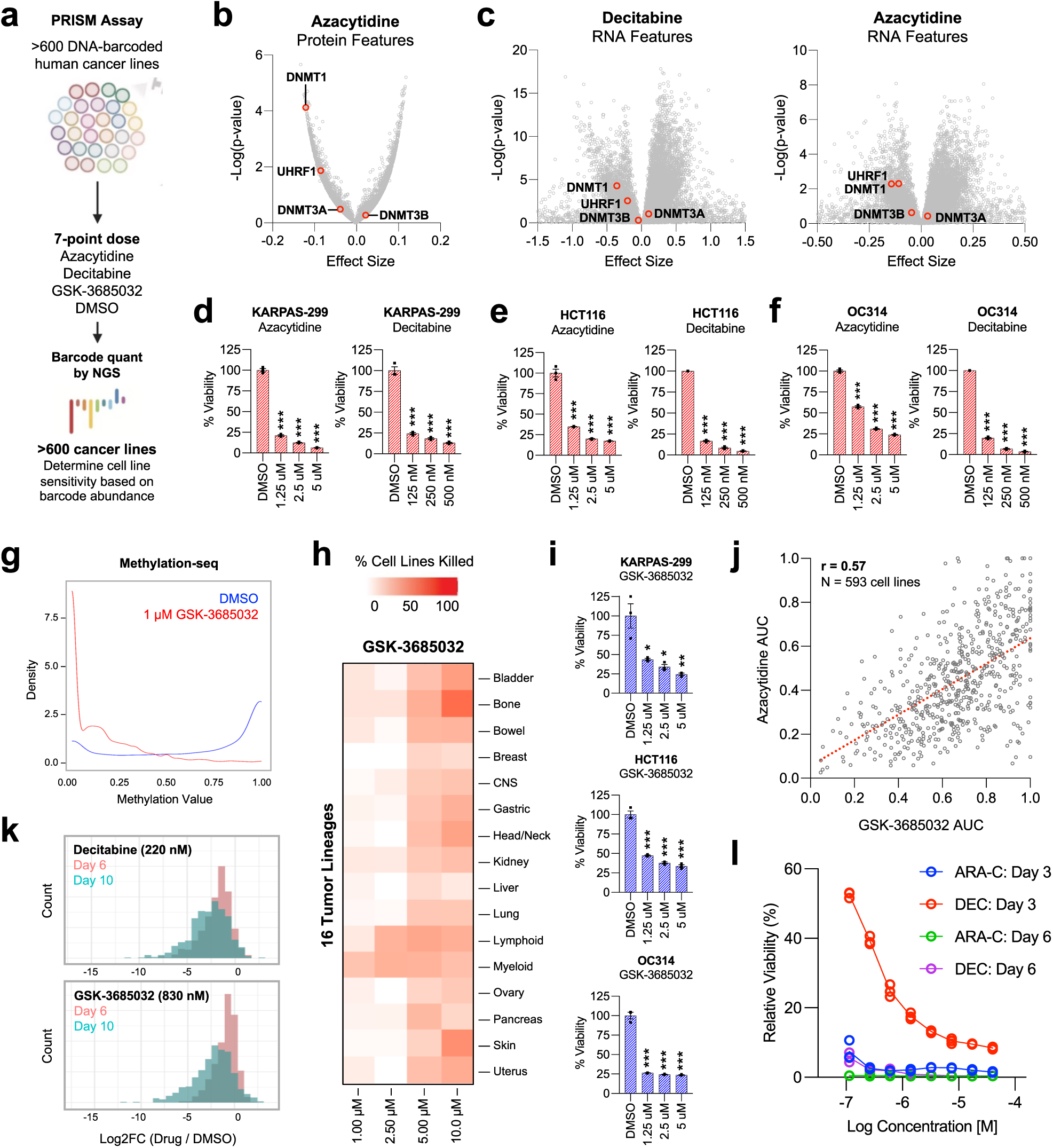
PRISM screens with AZA, DEC, and GSK-3685032. **a.** PRISM screen: >600 barcoded human cancer cell lines were pooled and treated with AZA (n=3), DEC (n=3), or GSK-3685032 (n=3) in a 7-point dose response (0.25-20 µM for AZA, 0.05-5 µM for DEC, and 0.25-20 µM for GSK-3685032). Viability for each cell line was determined based on relative barcode abundance with drug treatment compared to a DMSO control. **b.** Analysis of protein expression features that predict sensitivity to AZA. **c.** Analysis of RNA expression features that predict sensitivity to DEC or AZA. **d-f.** Viability of KARPAS-299 (lymphoid), HCT116 (colorectal), and OC314 (ovarian) cell lines following 6-day treatment with AZA (n=3) or DEC (n=3), compared to DMSO (n=3). **g.** Histograms of genome-wide cytosine methylation values in MOLM-13 cells after 3-day treatment with 1 µM GSK-3685032 (n=1) or DMSO (n=1). **h.** Lineage analysis of GSK-3685032 killing activity in the PRISM screen. Heatmap indicates the percentage of sensitive cell lines (Log2FC<-2) for each lineage and drug dose. **i.** Viability of KARPAS-299, HCT116, and OC314 cell lines following 6-day treatment with GSK-3685032 (n=3) compared to DMSO (n=3). **j.** AUC correlation analysis of AZA and GSK-3685032 across the cancer cell lines in PRISM. **k.** Viability histograms for >400 cell lines treated with 220 nM DEC (n=3) (upper panel) or 830 nM GSK-3685032 (n=3) (lower panel) for 6 or 10 days in an extended timecourse PRISM screen. **l.** Relative viability of MOLM-13 cells treated with ARA-C (n=3) or DEC (n=3) across a dose range for 3 or 6 days. ^∗^p < 0.05, ^∗∗^p < 0.01, ^∗∗∗^p < 0.001, determined by a two-sided Student’s t test (compared to the DMSO control). Mean values are shown unless otherwise specified, and error bars represent ± SEM.

**Extended Data Figure 2:**
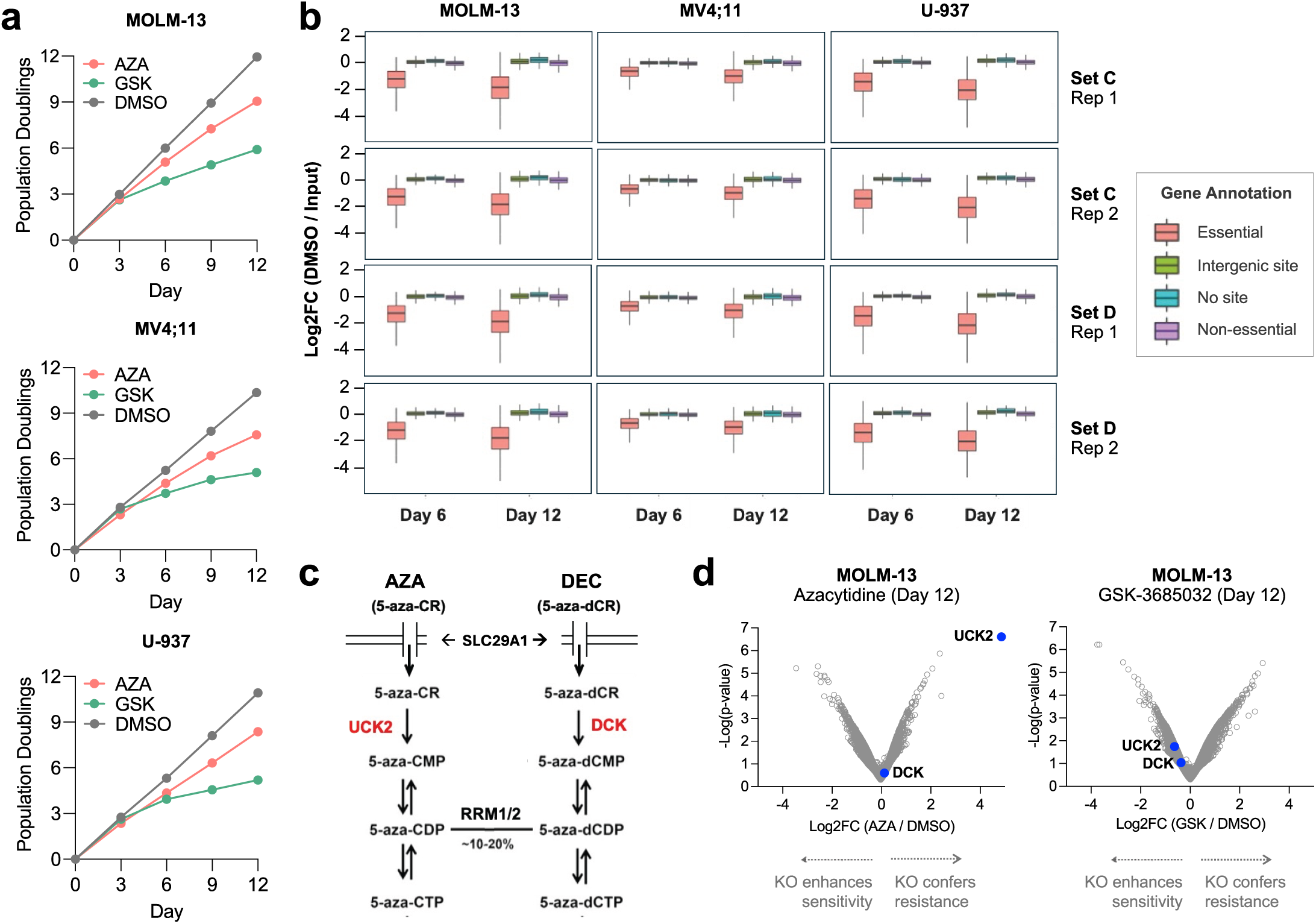
CRISPR drug modifier screens with AZA and GSK-3685032. **a.** Intra-screen growth curves of MOLM-13, MV4;11, or U-937 AML cell lines treated with AZA, GSK-3685032, or DMSO. **b.** Activity of sgRNAs predicted to target essential genes, intergenic sites (control), no genomic sites (control), or non-essential genes for each cell line (MOLM-13, MV4;11, or U-937), timepoint (day 6 or 12), sgRNA library set (set C or D), and replicate (replicate 1 or 2). Across all screens, sgRNAs targeting essential genes were depleted more than the control or non-essential sgRNA classes. **c.** Biochemical pathway for AZA and DEC cellular uptake and bioactivation. AZA and DEC enter the cell via the equilibrative nucleoside transporter SLC29A1 and are phosphorylated by UCK2 or DCK, respectively. 5-aza-CDP is converted to a deoxyribose form (5-aza-dCDP) by ribonucleotide reductase (RRM1/2), thereby enabling DNA incorporation. **d.** Volcano plots of MOLM-13 CRISPR drug modifier screens. Genetic deletion of UCK2 (but not DCK) rescues killing by AZA. Neither UCK2 knockout nor DCK knockout rescues killing by GSK-3685032.

**Extended Data Figure 3:**
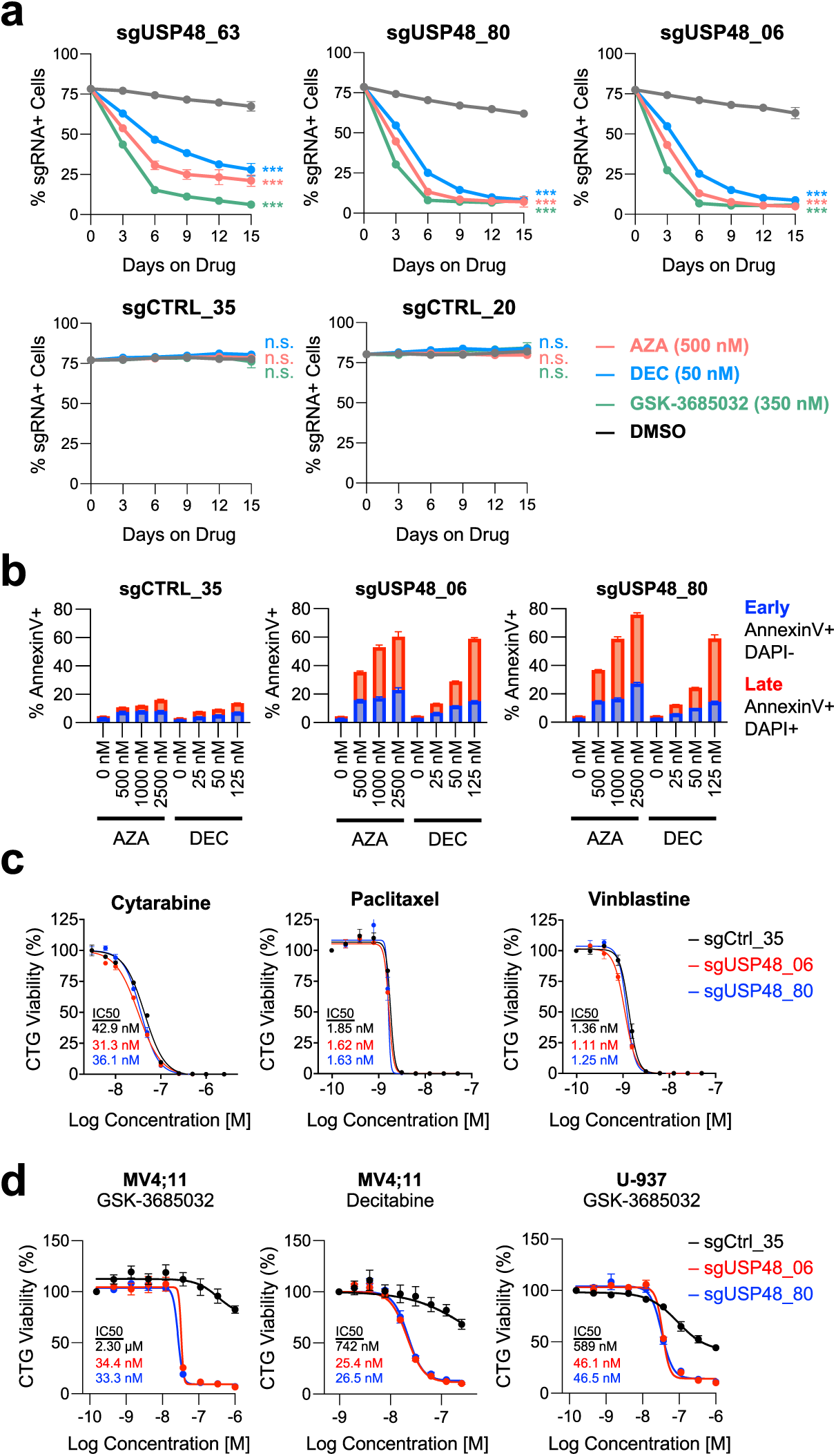
USP48 loss sensitizes cancer cells to DNMT1 inhibitor-induced cell death. **a.** Competition assays with MOLM-13 cells expressing sgRNAs targeting *USP48* (or control sgRNAs) after treatment with 500 nM AZA (n=3), 50 nM DEC (n=3), 350 nM GSK-3685032 (n=3), or DMSO (n=3). **b.** Annexin-V flow cytometry assays in *USP48*-intact (sgCTRL_35) or *USP48*-deleted (sgUSP48_06 or sgUSP48_80) MOLM-13 cells after 48-hr treatment with AZA (n=3), DEC (n=3), or DMSO (n=3). **c.** Dose-response curves of *USP48*-intact (sgCTRL_35) or *USP48*-deleted (sgUSP48_06 or sgUSP48_80) MOLM-13 cells after 72-hr treatment with chemotherapeutic agents (cytarabine, paclitaxel, or vinblastine) (n=3). **d.** Dose-response curves with *USP48*-intact (sgCTRL_35) or *USP48*-deleted (sgUSP48_06 or sgUSP48_80) AML cell lines treated with DEC (n=3) or GSK-3685032 (n=3) for 72 hrs. ^∗^p < 0.05, ^∗∗^p < 0.01, ^∗∗∗^p < 0.001, determined by an unpaired, two-sided Student’s t-test. Mean values are shown unless otherwise specified, and error bars represent ± SEM.

**Extended Data Figure 4:**
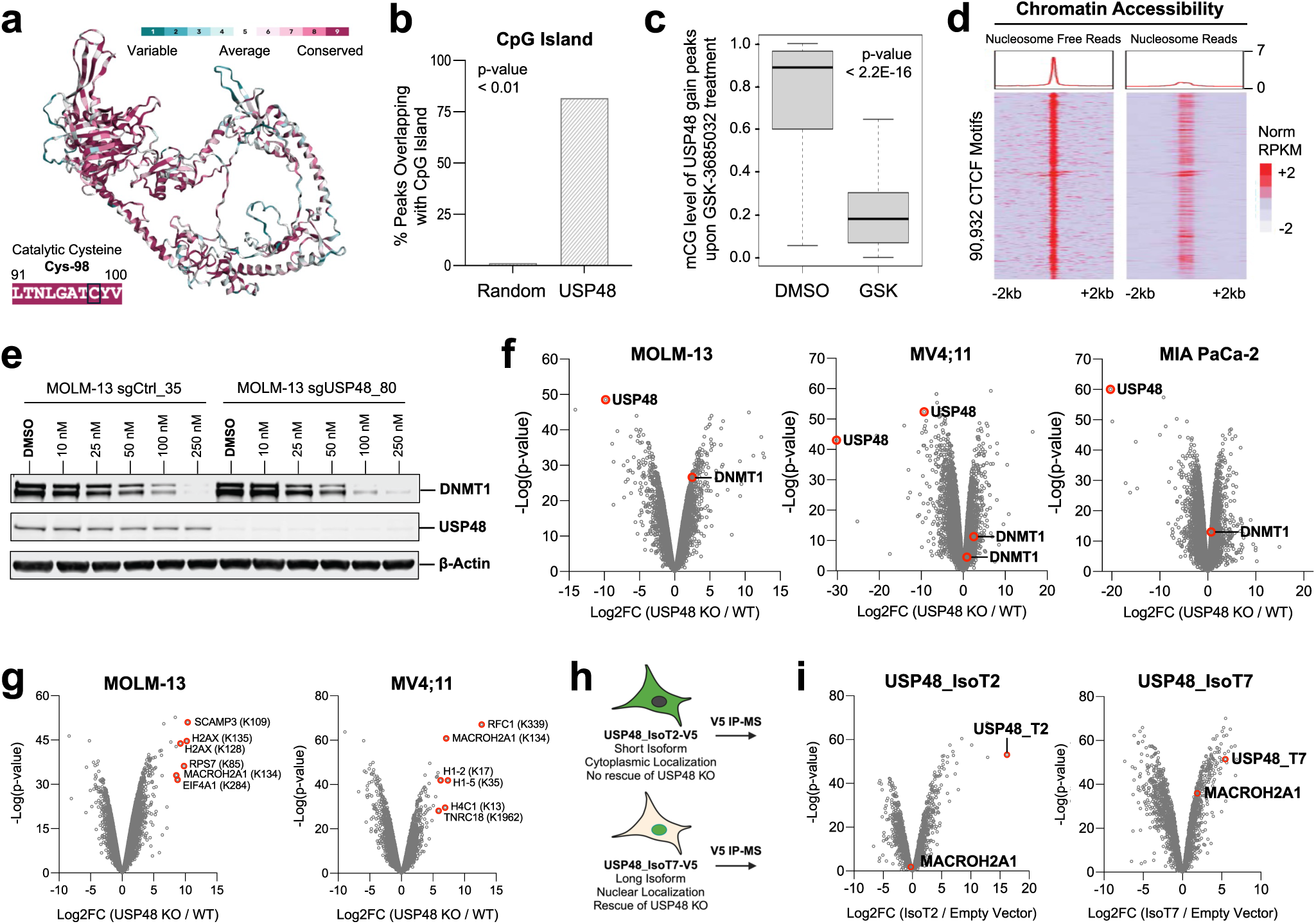
USP48 is a catalytically-active DUB that targets variant histones. **a.** ConSurf amino acid residue conservation score projected onto the AF2 structural model of USP48. The peptidase domain and catalytic cysteine (Cys-98) of USP48 are highly conserved from yeast to humans. **b.** CpG island enrichment analysis of USP48 genomic binding sites in MOLM-13 cells. **c.** Methyl-seq analysis of novel USP48 genomic binding sites gained in GSK-3685032-treated MOLM-13 cells. **d.** CTCF motif analysis of the MOLM-13 ATAC-seq profiles. **e.** Western blot analysis of DNMT1 protein levels in isogenic *USP48*-intact (sgCTRL_35) and *USP48*-deleted (sgUSP48_80) MOLM-13 cells after 48-hr treatment with GSK-3685032. **f.** Whole proteome profiling of *USP48*-deleted (sgUSP48_06) cells compared to *USP48*-intact (sgCTRL_35) cells for MOLM-13 (n=2), MV4;11 (n=3), or MIA PaCa-2 (n=3), highlighting preserved DNMT1 protein stability in the absence of *USP48*. **g.** Ubiquitylome (normalized to proteome) profiling in MOLM-13 (n=2) or MV4;11 (n=3) following *USP48* deletion (sgUSP48_06) compared to intact *USP48* (sgCTRL_35). **h.** IP-MS strategy to identify chromatin-based interacting partners of USP48 using distinct USP48 isoforms. The long isoform (IsoT7) of USP48 is a nuclear protein, while the short isoform (IsoT2) is a cytoplasmic protein and can be used as a control. **i.** IP-MS volcano plots of the interacting partners of the short isoform (n=3) or long isoform (n=3) of USP48. MACROH2A1 selectively binds the USP48 long isoform.

**Extended Data Figure 5:**
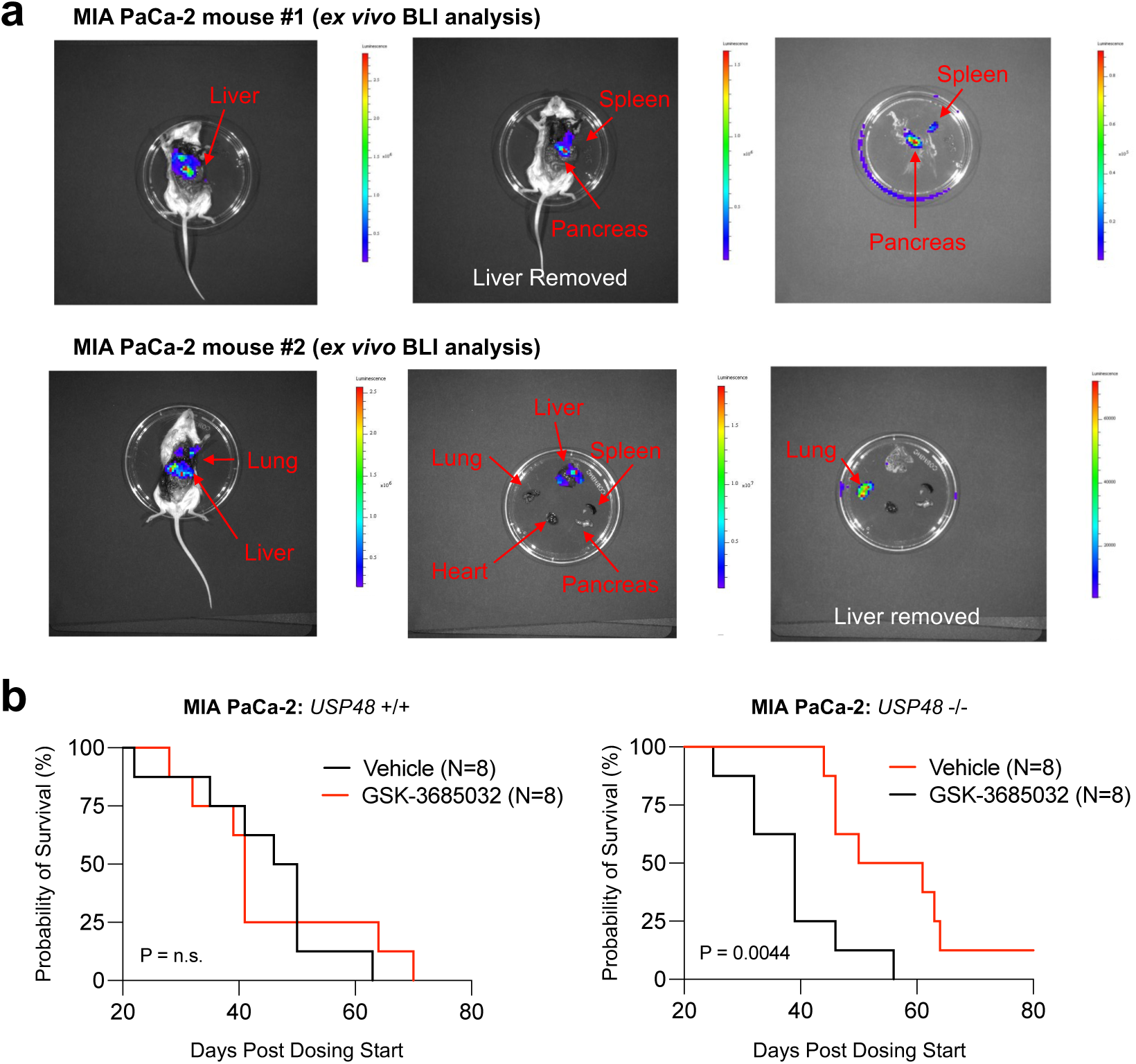
*In vivo* metastatic pancreatic cancer model. **a.** *Ex vivo* BLI reveals extensive metastatic tumor formation in NSG mice following intracardiac injection of MIA PaCa-2 pancreatic cancer cells. **b.** Kaplan-Meier survival analysis of *USP48* +/+ or *USP48* −/− MIA PaCa-2 metastatic tumor cohorts treated with vehicle or GSK-3685032 (30 mg/kg BID).

