## Supplementary Information for "Discovery of chromatin-based determinants of azacytidine and decitabine anti-cancer activity"

### SUPPLEMENTARY METHODS

#### Cell Culture

MOLM-13, MV4;11, U-937, OCI-AML2, and KARPAS-299 cell lines were cultured in suspension in RPMI-1640 (Gibco) supplemented with 20% fetal bovine serum (FBS) (Sigma-Aldrich, F7524) and 1% penicillin/streptomycin (P/S) (Gibco, 15140122). The isogenic *TP53* wildtype and *TP53* deleted MOLM-13 cell lines (gifts from Ben Ebert) were cultured similarly to the parental MOLM-13 cell line[1]. MIA PaCa-2, HCT116, OC314, NCIH-1299, and KNS60 cells were cultured in RPMI-1640 with 10% FBS and 1% P/S. HEK293T cells were maintained in DMEM (Gibco) with 10% FBS and 1% P/S. Prior to all CRISPR screens, cell lines were STR profiled and tested for mycoplasma through Genetica Cell Line Testing (LabCorp). Puromycin concentrations for selection were as follows: 4 µg/mL (MOLM-13), 2 µg/mL (MV4;11, U-937, OCI-AML2, and KARPAS-299), or 1 µg/mL (MIA PaCa-2 and HEK293T).

#### Compound Treatments

The following compounds were obtained commercially, resuspended in DMSO at a concentration of 10-40 mM, and stored in aliquots at -80°C for long term use: Azacytidine (Selleck Chemicals, S1782), Decitabine (Selleck Chemicals, S1200), GSK-3685032 (MedChem Express, HY-139664), (S)-GSK-3685032 (MedChem Express, HY-139664B), (R)-GSK-3685032 (MedChem Express, HY-139664A), GSK-3484862 (MedChem Express, HY-135146), Paclitaxel (Selleck Chemicals, S1150), Doxorubicin (Selleck Chemicals, S1208), Cytarabine (Selleck Chemicals, S1648), Vinblastine (Selleck Chemicals, S4505), PR-619 (MedChem Express, HY-13814), 17e (Naumann Lab, custom synthesis)[2], and dTag$^{V}$-1 (Tocris, 6914). For drug treatments across a multi-point dose range or for treatments in multi-well plates, compounds were added using the D300e Digital Dispenser (Tecan, 30100152). DMSO volume was normalized across control and treatment conditions for all experiments.

#### PRISM Viability Screening

Extended PRISM assay: The extended 10-day PRISM assay incorporated several modifications compared to the 6-day assay: (1) Compounds were pre-dispensed to 24-well poly-D-lysine-coated plates, which were stored at -20°C and thawed at 37°C for 2 hrs before cell seeding directly onto the assay-ready plates; (2) A revised cell pooling strategy utilized 475 PRISM cell lines distributed across 14 subpools with seeding densities optimized for a 10-day timepoint; (3) Duplicate plates were prepared with a set lysed at day 6 and the second set at day 10 with media and compounds refreshed at day 6.

CRISPR/Cas9 PRISM assay: For evaluating the combination effect of *USP48* knockout and GSK-3685032 treatment, 483 PRISM cell lines distributed across 5 subpools were transduced by spinfection with the pXPR_044 all-in-one vector for co-expression of *Cas9* and an sgRNA targeting either *USP48* (sgUSP48_80) or an intergenic control sequence (sgCTRL_00). PRISM subpools were resuspended at a concentration of 3E6 cells/mL in RPMI-1640 without phenol red supplemented with 10% FBS, 1% P/S, and 10 µg/mL polybrene. Equal volumes (1 mL) of cell suspension and lentivirus were combined and plated in individual wells on a 6-well plate. The plate was centrifuged at 2000 RPM at 37°C for 2 hrs followed by the addition of 1 mL fresh media to each well. After overnight incubation, subpools were combined and expanded in T-175 flasks for 72 hrs followed by isolation of mCherry-positive cells by FACS, yielding

*USP48* wildtype and *USP48* knockout PRISM pools. Each sorted PRISM pool was plated in 6-well dishes, allowed to recover for 72 hrs, and then treated with GSK-3685032 (0.1, 0.5, or 2.5 µM) or DMSO for 72 hrs followed by cell lysis.

Library preparation and next generation sequencing: Matched treatment conditions (same compound dose, timepoint, and replicate) were collapsed to single lysates prior to PCR, with hematopoietic cell line pools processed separately from solid tumor cell line pools. Collapsed lysates were denatured at 95°C for 5 min and then equal lysate volumes from each well were added to a custom PCR master mix for PRISM cell barcode amplification. Ten unique control barcodes were spiked into each well before PCR to account for variation in amplification and sequencing across treatment conditions. PCR products underwent quality control via gel electrophoresis (Invitrogen, A42100) to confirm successful barcode amplification. Libraries were prepared by collapsing and purifying PCR products using the ThermoFisher PureLink PCR Purification Kit (ThermoFisher, K310001), according to the manufacturer's protocol. The PCR products were quantified via the Qubit High Sensitivity Assay (ThermoFisher, Q33231) and TapeStation, and submitted for next generation sequencing on the Illumina NextSeq platform.

Data analysis: PRISM screen data analysis followed the SUSHI pipeline for DNA sequencing-based screens (https://github.com/cmap/sushi). Cell line barcodes were extracted from FASTQ reads and tallied to create raw count tables. Counts in each PCR well were log10 transformed and mapped to log10 concentrations using the set of 10 control barcodes spiked in at different concentrations. To calculate the fold change in viability, the normalized counts for each treatment condition were divided by the normalized counts of the corresponding collapsed control for each biological replicate. These values were log2 transformed, and the median of the biological replicates was reported as the final log2 fold change (Log2FC) value. In each treatment or control condition, a cell line was removed if it had a standard error (MAD/sqrt(n), where MAD is the median absolute deviation and n is the number of replicates) larger than 0.5 across its biological replicates or corresponding negative controls. Area under the curve (AUC) values for each cell line were calculated for treatment conditions as the arithmetic mean of the fold change viability values (capped at a maximum of 1) across doses (minimum of 3 doses required). Simple linear regression coefficients were calculated for each AUC or dose-level Log2FC profile against mRNA and protein expression features (DepMap 23Q2 Release) using the custom analysis tool (https://depmap.org/portal/interactive/custom_analysis) from the Broad Institute's DepMap portal[3]. Calculated effect sizes and significance values (adjusted p-values) are reported, as described previously[4]. For the AUC correlation analysis for AZA, DEC, and GSK-3685032, the correlation coefficient (r) was calculated based on a simple linear regression model using GraphPad PRISM software. For the lineage sensitivity analysis, the percentage of sensitive (Log2FC<-2) cell lines within each lineage for each drug dose is reported; lineages with fewer than 10 cell lines represented were excluded from the analysis.

**Humagne CRISPR/Cas12a Drug Modifier Screens**

Data analysis: PoolQ (version 3.05) was used to deconvolute sgRNA sequences and generate read count matrices. From the input condition, sgRNA sequences with read counts under 50 were identified and removed across all conditions. Next, log2 TPM values were calculated for each condition. From log2 TPM values, L2FC values were computed for each treatment relative to DMSO at each time point and centered such that the mean L2FC of the control sgRNAs corresponded to zero. Quality checks included verifying high replicate concordance and strong drop-out of sgRNAs targeting common essential genes. Gene-level

L2FC values were calculated by taking the average across all sgRNAs targeting the same gene; p-values were calculated on the GPP Web Portal (https://portals.broadinstitute.org/gpp/public/) using the hypergeometric analysis method, where sgRNAs are ranked and assigned p-values using a hypergeometric distribution. For screens with more than 2 timepoints, Chronos (version 1) was used to generate gene effect scores, as previously described[5]. The top-scoring sensitizer genes were defined by selecting genes with an average L2FC<-2 for the day 6 and 12 timepoints for AZA and an average L2FC<-2 for the day 6 and 12 timepoints for GSK-3685032. Z-scored L2FC values were also calculated for each treatment condition.

The codependency analysis was performed using CRISPR gene effects from DepMap version 2023Q2 (https://depmap.org). Pairwise Pearson correlations were calculated for every unique gene pair using the gene effect scores across all DepMap cell lines. The heatmap was generated using the R package ComplexHeatmap (version 2.22.0) with its default hierarchical clustering algorithm. For pathway enrichment analysis of the 37 top-scoring sensitizer genes, we performed gene set enrichment analysis (GSEA) using the canonical pathway (CP) reactome gene sets in the Molecular Signatures Database (MSigDB) public portal (https://www.gsea-msigdb.org/gsea/msigdb).

**Bison CRISPR/Cas9 Drug Modifier Screens**

Bison screens: The Bison CRISPR/Cas9 sgRNA library (Addgene, 169942) contains 2,852 sgRNAs targeting 713 E1, E2, and E3 ubiquitin ligases, deubiquitinases, and control genes. The library was cloned into the pXPR_003 vector, and lentivirus was produced as previously described[6]. Library transduction, drug treatments, and cell reseeding were performed similarly to the Humagne CRISPR screens. The following compounds and doses (in addition to a DMSO-only control) were used for the KARPAS-299 and MIA PaCa-2 screens: 0.2 µM DEC and 2 µM GSK-3685032. Cell pellets were resuspended in 100 µL direct lysis buffer (1 mM CaCl2, 3 mM MgCl2, 1 mM EDTA, 1% Triton X-100, Tris pH 7.5) with freshly supplemented 0.2 mg/mL proteinase K (Qiagen, 19133). Lysates were incubated at 65°C for 15 min followed by 95°C for 10 min. For library amplification, 25 µL of lysate for each sample was used in a 50 µL reaction volume with 0.04 U Titanium Taq (Takara, 639210), 0.5x Titanium Taq buffer, 800 µM dNTP mix, 200 nM P5-SBS3-Stagger-pXPR003 forward primer, and 200 nM P7-Barcode-SBS12-pXPR003 reverse primer with the following heat cycle: 94°C for 5 min, 32 cycles of [94°C for 30 s, 58°C for 15 s, 72°C for 30 s], and 72°C for 2 min. An equal amount of all samples was pooled and subjected to preparative agarose electrophoresis followed by gel purification (Qiagen, 28704). Eluted DNA was further purified by AMPure XP Bead (Beckman Coulter, A63881). Amplified sgRNAs were quantified using the Illumina NextSeq platform (Genomics Platform, Broad Institute).

Data analysis: The data analysis pipeline consisted of the following steps: (1) Normalize each sample to the total number of reads; (2) For each sgRNA, calculate the ratio of reads in the drug treatment samples (DEC or GSK-3685032) versus the corresponding DMSO control samples and rank the sgRNAs; (3) Sum the ranks for each sgRNA across all replicates; (4) Determine the gene rank as the median rank of the 4 guides targeting it; (5) Calculate p-values by simulating a distribution with sgRNAs that have randomly assigned ranks over 100 iterations. The R scripts for these steps were previously published[6].

**Cas9/sgRNA Cell Competition Assays**

Control sgRNAs (sgCTRL_00, sgCTRL_35, or sgCTRL_20) and *USP48* sgRNAs (sgUSP48_06, sgUSP48_80, or sgUSP48_63) were cloned into a modified version of pLentiGuide-Puro (Addgene, 52963). The modified vector consists of a U6 promoter driving sgRNA expression and an EF1α promoter driving expression of a fluorescent protein CDS (*RFP657* or *BFP*) followed by a P2A sequence and *PAC* (puromycin resistance). Cas9-expressing AML cell lines were transduced with a control sgRNA (sgCTRL_00, sgCTRL_35, or sgCTRL_20) with co-expression of *BFP-P2A-PAC* and selected with puromycin (4 µg/mL), resulting in 3 BFP+ *USP48* wildtype lines. Separately, Cas9-expressing AML cell lines were transduced with a *USP48* sgRNA (sgUSP48_06, sgUSP48_80, or sgUSP48_63) with co-expression of *RFP657-P2A-PAC* and selected with puromycin, resulting in 3 RFP657+ *USP48* knockout lines. Each wildtype line (BFP+) was mixed with a knockout line (RFP657+) at a ratio of 20:80, plated in 12-well dishes, and treated with DMSO or 0.5 µM GSK-3685032 with drug refreshed every 2 days. The percentage of BFP+ and RFP+ cells was measured by flow cytometry over a 10-day period on the Cytoflex LX (Beckman Coulter). A reduction in the percentage of RFP657+ cells over time reflects decreased viability of *USP48* knockout cells compared to *USP48* wildtype cells.

**Immunoblotting**

Western blots were performed using a standard protocol. Briefly, cells were washed with chilled PBS and resuspended in RIPA buffer (ThermoFisher) with Halt protease and phosphatase inhibitor cocktail (ThermoFisher, PI78441). Lysates were spun down to remove debris, diluted 2-fold in Laemmli sample buffer, boiled at 100°C for 5 min, and immediately run on NuPAGE Bis-Tris precast protein gels (ThermoFisher, WBT41212). Transfers were performed using Immobilin-P membranes (Millipore) (100 V for 60 min). Transfer membranes were blocked in TBS with 5% BSA for 1 hr at room temperature, stained with primary antibody (1:1000) overnight at 4°C, washed 5 times with TBS-T, stained with secondary antibody (1:5000) for 1 hr at room temperature, washed 5 times with TBS-T, and rinsed twice with TBS. The Licor system and Odyssey CLx imager were used to visualize the blot. The following antibodies were commercially-obtained for western blotting: USP48 (Abcam, ab72226), DNMT1 (D63A6) (Cell Signaling Technology, 5032), Phospho Histone H2A.X (Ser139) (20E3) (Cell Signaling Technology, 9718), V5-tag (D3H8Q) (Cell Signaling Technology, 13202), P53 (DO-1) (Cell Signaling Technology, 18032), P21 (12D1) (Cell Signaling Technology, 2947), GAPDH (14C10) (Cell Signaling Technology, 2118), Ubiquitin (Santa Cruz, sc-8017), TUBB (Cell Signaling Technology, 2146), and ACTB (8H10D10) (Cell Signaling Technology, 3700).

**5-methylcytosine Dot Blot**

Parental MOLM-13 cells were plated at 200,000 cells/mL in 12-well dishes and treated with AZA or GSK-3685032 in a dose response (0, 0.1, 0.25, 0.5, 1, or 2.5 µM) for 72 hrs with daily replenishment of drug. Following drug treatment, genomic DNA (gDNA) was isolated from cell pellets using the QIAamp Blood & Cell Culture DNA Mini Kit (Qiagen, 51304). The gDNA concentration was measured with the Qubit dsDNA Quantitation (Broad Range) Assay Kit (Thermo Fisher Scientific, Q32850) and was normalized across all samples with the addition of ddH2O. Following treatment with RNAse A, samples were denatured at 95°C for 10 min and then cooled on ice for 5 min. A total of 100 ng of DNA for each sample was loaded directly onto a nitrocellulose membrane and air-dried for 5 min. The membrane was then blocked in TBS with 5% BSA for 1 hr at room temperature, incubated with anti-5-methylcytosine (5-mC) antibody (Zymo, A3002) (1:1000) at 4°C overnight, washed 3 times with TBS-T, stained with anti-mouse IgG IRDye 680RD

(LI-COR, 926-68070) for 90 min at room temperature, washed 3 times with TBS-T, and imaged on the Odyssey CLx imager.

#### *In Vitro* DUB Assay

Recombinant His6-USP48-FL (Boston Biochem, E-614-050) was incubated for 5 min at 30°C with continuous shaking in buffer (50 mM HEPES pH 7.5, 100 mM NaCl) supplemented with dithiothreitol (DTT, 1 mM) (AppliChem, A2948) and 4-(2-aminoethyl)benzolsulfonylfluorid (AEBSF, 1 mM) (AppliChem, A1421) in the presence or absence of either PR-619 or 17e DUB inhibitors. After the addition of Ub4 (K63) (LifeSensors, SI6304), the samples were incubated at 30°C for up to 60 min with constant shaking. Samples were then boiled (5 min, 95°C) with 2x Laemmli sample buffer and β-mercaptoethanol (Sigma Aldrich, 63689), separated by SDS-PAGE, and analyzed by immunoblot with antibodies against Ubiquitin (Santa Cruz, sc-8017) and USP48 (Abcam, ab72226).

#### Conditional Degradation of USP48

The *USP48_IsoT7* (with C-terminal V5 tag) CDS was cloned into the pDeg vector (Addgene, 185760-185779), as previously described[7]. We generated a pDeg vector set containing distinct degron tags (dTAG, HaloTag, or Ikzf3b), degron tag placements (N-terminal or C-terminal), and promoters (PGK or SFFV) driving expression of the *USP48_IsoT7* CDS. Each vector was tested in HEK293T cells to ensure physiologic USP48 expression levels and efficient USP48 degradation by western blot after adding the respective degron compound for 24 hrs. The vector with an SFFV promoter and C-terminal dTAG was chosen for follow-up experiments in MOLM-13 cells. *USP48* knockout MOLM-13 cells were stably transduced with the *USP48* C-terminal dTAG vector (USP48-dTAG add-back) followed by selection with puromycin (4 µg/mL). The USP48-dTAG add-back cells were then assessed for USP48-dTAG degradation after treatment with $dTAG^{V}$-1 (Tocris, 6914). For viability experiments, the USP48-dTAG add-back cells were pre-treated with 1 µM GSK-3685032 for 3 days followed by drug washout, exposed to DMSO or $dTAG^{V}$-1 (0.5, 1, or 2 µM) to induce USP48 degradation, and counted for viable cells approximately every 3 days over a 2-week timecourse.

#### *USP48* Rescue Experiments

We created a modified version of the pLX_203 vector (Broad GPP) with a PGK promoter driving expression of *PAC* (puromycin resistance) followed by a P2A sequence and *RFP657* and a separate EF1α promoter driving expression of a codon-optimized version of the *USP48* CDS (*WT_IsoT2*, *WT_IsoT7*, or *C98S_IsoT7*) with a C-terminal V5 tag. Given the presence of silent point mutations in the CDS, these *USP48* cDNAs were resistant to cutting by sgUSP48_80. Using these constructs, MIA PaCa-2 cells were transduced with the *USP48* CDS (*WT_IsoT2*, *WT_IsoT7*, or *C98S_IsoT7*) or an empty vector control and selected with puromycin (1 µg/mL), yielding 3 isogenic *USP48* overexpression lines and 1 control line. Each of these isogenic lines was lentivirally-transduced with a *USP48* sgRNA (sgUSP48_80) or control sgRNA (sgCTRL_35) with co-expression of *GFP*, expanded for 3 days, and then treated with DMSO or 0.5 µM GSK-3685032 with drug refreshed every 3 days. The percentage of GFP+ cells was measured over a 13-day timecourse by flow cytometry on the Cytoflex LX (Beckman Coulter). A reduction in the percentage of GFP+ cells over time with sgUSP48_80 (compared to sgCTRL_35) indicates that the *USP48* CDS construct did not effectively rescue *USP48* knockout.

### Immunofluorescence

MIA PaCa-2 cells were lentivirally-transduced with the *USP48* CDS (*WT_IsoT2* or *WT_IsoT7*) with a C-terminal V5 tag or empty vector and plated on a 96-well PhenoPlate (Perkin Elmer). The next day, wells were washed with PBS, fixed with 4% paraformaldehyde (PFA) for 15 min at room temperature, washed twice with PBS, permeabilized with 0.5% Triton X-100 for 5 min at room temperature, washed twice with PBS, blocked in Cell Signaling Blocking Buffer (Cell Signaling Technology, 12411) for 1 hr at room temperature, and incubated at 4°C overnight with anti-V5-tag primary antibody (D3H8Q) (Cell Signaling Technology, 13202) at a 1:1000 concentration in Immunofluorescence Antibody Dilution Buffer (Cell Signaling Technology, 12378). The next day, wells were washed 3 times with PBS, incubated with secondary antibody conjugated to Alexa-488 (Invitrogen, A11034) at a 1:500 concentration in Immunofluorescence Antibody Dilution Buffer for 1 hr in the dark at room temperature, and washed 3 times with PBS. DAPI was added immediately prior to imaging on the Phenix High-Content Imaging System (Revvity) and analyzed with the Harmony High-Content Imaging and Analysis Software (Revvity).

### Methylation-seq

All Methyl-seq enzymatic treatment steps, library preparation, and next generation sequencing were performed by Azenta (South Plainfield, NJ, USA).

<u>Enzymatic Methyl-seq</u>: Each gDNA sample was initially treated with RNase (New England Biolabs) to remove RNA contaminants. The DNA sequencing library was prepared using 100 ng gDNA combined with 0.001 ng CpG methylated pUC19 and 0.02 ng unmethylated lambda control DNA and sheared using the Covaris LE220 Focused-ultrasonicator for an average 350 bp size. The sheared material was transferred to a PCR strip tube to begin library construction. NEBNext Enzymatic Methyl-Seq Kit (New England Biolabs) was used according to the manufacturer's instructions for end repair, A-tailing, and adaptor ligation of EM-seq adaptor. The ligated samples were cleaned up according to the manufacturer's protocol with NEBNext Sample Purification Beads. Ligated DNA was oxidized by TET2 enzymatic reaction, which was initiated by adding Fe (II) solution and then incubated for 1 hr at 37°C. Following this step, Stop Reagent was added and incubated for 30 min at 37°C. Oxidized DNA was cleaned up with NEBNext Sample Purification Beads, denatured by formamide, and deaminated with an APOBEC enzymatic reaction, which was incubated at 37°C for 3 hrs.

<u>Library preparation and sequencing</u>: Deaminated DNA was cleaned up with NEBNext Sample Purification Beads, and then PCR amplified with NEBNext Q5U Master Mix and EM-Seq Index Primers. Amplified library was cleaned up with NEBNext Sample Purification Beads. The final library was assessed with Qubit 4.0 Fluorometer (ThermoFisher Scientific) and Agilent TapeStation (Agilent Technologies), and quantified by qPCR (KAPA Biosystems). The sequencing libraries were multiplexed and sequenced via Illumina NovaSeq using a 2x150 paired end (PE) configuration. Image analysis and base calling were conducted by the control software. Raw sequence data (.bcl files) generated from the sequencer were converted into fastq files and de-multiplexed using Illumina's bcl2fastq 2.17 software. One mismatch was allowed for index sequence identification.

Data analysis: Whole-genome DNA methylation data were analyzed using the methylCtools pipeline (https://github.com/hovestadt/methylCtools/tree/master). The analysis began by converting the reference genome (hg38) into a C-to-T-converted reference file (hg38.conv.fa) using the methylCtools faconv command. The converted reference genome was then indexed with the bwa index command to generate the hg38.conv.bwt file required for subsequent alignments. Additionally, genomic position information was extracted from the reference genome using the methylCtools fapos command, resulting in the hg38.pos file. Paired-end sequencing reads (reads1.fq and reads2.fq) underwent C-to-T conversion using the methylCtools fqconv command, generating reads1.conv.fq and reads2.conv.fq. These converted reads were aligned separately to the converted reference genome using the bwa aln command, producing reads1.conv.sai and reads2.conv.sai files. Paired-end alignments were then combined and converted into BAM format using the bwa sampe and samtools view commands, resulting in reads.conv.bam. The aligned reads were further processed using the methylCtools bconv command and sorted with samtools sort, yielding reads.bam. Duplicate reads were identified and marked using the Picard MarkDuplicates tool, resulting in the final aligned file (sample.bam). Methylation calling was performed using the methylCtools bcall command to generate the final methylation call file (sample.call).

**ATAC-seq**

All ATAC-seq sample processing steps, Tn5 treatment, library preparation, and next generation sequencing were performed by Azenta (South Plainfield, NJ, USA).

Tn5 treatment: Live cell samples were thawed, washed, and treated with DNAse I (Life Tech, EN0521) to remove genomic DNA contamination. The samples were assessed for cell viability using the Countess Automated Cell Counter (ThermoFisher Scientific). After cell lysis and cytosol removal, nuclei were treated with Tn5 enzyme (Illumina, 20034197) for 30 min at 37°C and purified with the Minelute PCR Purification Kit (Qiagen, 28004) to produce tagmented DNA samples.

Library preparation and sequencing: Tagmented DNA was barcoded with Nextera Index Kit v2 (Illumina, FC-131-2001) and amplified via PCR prior to a SPRI bead cleanup to yield purified DNA libraries. The sequencing libraries were multiplexed and sequenced via Illumina NovaSeq using a 2x150 paired end (PE) configuration. Image analysis and base calling were conducted by the Control software. Raw sequence data (.bcl files) generated from the sequencer were converted into fastq files and de-multiplexed using Illumina's bcl2fastq 2.17 software. One mismatch was allowed for index sequence identification.

Data analysis: Sequencing adapters and low-quality bases were trimmed using trim galore. Cleaned reads were then aligned to hg38 using bowtie2. Aligned reads were filtered using samtools 1.9 to keep alignments that are aligned concordantly and are the primary called alignments. PCR or optical duplicates were marked using Picard and removed. Prior to peak calling, reads mapping to the mitochondrial (mt) genome were called and filtered, and reads mapping to unplaced contigs were removed. MACS2 was used for peak calling to identify open chromatin regions..

**ChIP-seq**

All of the ChIP sample processing steps, library preparation, and next generation sequencing were performed as previously described[8].

Double cross-linking: Briefly, 20E6 cells were collected by centrifugation at 1200 RPM, washed twice with cold PBS and resuspended in 2 mL of room temperature PBS. Next, 8 µL of 0.5 M disuccinimidyl glutarate (DSG) (ThermoFisher, 20593) was added, and samples were incubated for 25 min at room temperature with gentle rocking. Cells were pelleted by centrifugation at 1600 RPM and resuspended in 3.32 mL of PBS and 3.32 mL of 2% formaldehyde and incubated for 15 min at room temperature in the dark with gentle rocking. The cross-linking reaction was quenched with the addition of 666 µL of 1 M Tris-HCl pH 8.0 and 333 µL of 2.5 M glycine. Cells were pelleted by centrifugation at 700 × g and washed twice with room-temperature PBS.

Nuclear isolation: Cross-linked cells were resuspended in LB1 buffer (50 mM HEPES-KOH pH 7.5, 140 mM NaCl, 1 mM EDTA, 10% Glycerol, 0.5% NP-40, 0.25% Triton X-100, complete protease inhibitor) and rotated for 10 min at 4°C. Cells were pelleted by centrifugation at 1400 × g at 4°C and resuspended in LB2 buffer (10 mM Tris-HCl pH 8.0, 200 mM NaCl, 1 mM EDTA, 0.5 mM EGTA, complete protease inhibitor) and rotated for 10 min at 4°C. Cells were pelleted and resuspended in LB3 buffer (10 mM Tris-HCl pH 8.0, 100 mM NaCl, 1 mM EDTA, 0.5 mM EGTA, 0.1% Na-deoxycholate, 0.5% Na-Lauroylsarcosine, 1% Triton X-100, complete protease inhibitor) and sonicated for 50 min with Peak Power 140 Duty Factor 5.0 Cycles/Burst 200 (Covaris, E220).

Immunoprecipitation: Antibody-bead conjugates were prepared with either Protein-A or Protein-G Dynabeads (ThermoFisher). The ChIP-grade USP48 antibody was obtained commercially (Abcam, ab72226). Beads were washed twice with ChIP dilution buffer (0.01% SDS, 1.1% Triton X-100, 1.2 mM EDTA, 16.7 mM Tris HCl pH 8.0, 167 mM NaCl) and resuspended with 4.2 µL of beads/µg of antibody and rotated overnight at 4°C. Sonicated samples were transferred to DNALoBind tubes (Eppendorf) and clarified by centrifugation. Antibody-bead complexes were applied to cleared lysates and rotated overnight at 4°C. Beads were washed 6 times with RIPA buffer (50 mM Hepes-KOH pH 7.5, 500 mM LiCl, 1 mM EDTA, 0.7% Na-deoxycholate, 1% NP-40), twice with Buffer 500 (0.5 g deoxycholic acid, 1 mM EDTA, 5 mM Tris-HCl pH 8.1, 1% Triton X-100, and 0.02% NaN3), and twice with LiCl Buffer (2.5 g deoxycholic acid, 1 mM EDTA, 250 mM LiCl, 0.5% NP-40, 10mM Tris-HCl pH 8.0, 0.02% NaN3). Beads were washed briefly with TE buffer, 100 µL of ChIP elution buffer (50 mM Tris-HCl pH 8.0, 10 mM EDTA, 1% SDS in distilled water) was added to the beads, and samples were de-crosslinked overnight at 65°C. DNA was purified by column purification using DNA Clean and Concentration (Zymo, D4014).

Library preparation and sequencing: Library preparation was performed according to manufacturer's instructions using the ThruPlex DNA-Seq Kit (Takara, R400676) with 10 µL of sample. Fragment length and concentration were assessed using TapeStation (Agilent) and Qubit (ThermoFisher). Libraries were sequenced on an Illumina NextSeq 550 sequencer with a High Output Illumina Kit v2.5 (75 cycles) and using paired-end sequencing.

Data analysis: Paired-end reads were first trimmed using the Trim Galore (version 0.4.5) pipeline (https://github.com/FelixKrueger/TrimGalore), followed by read alignment to the hg38 genome using Bowtie 2 (Version 2.3.2) with the parameters: -t -q -N 1 -L 25 -X 700 no-mixed no-discordant. PCR duplicates were removed using "MarkDuplicates" from Picard Tools (version 2.8.0) (https://broadinstitute.github.io/picard/). Only non-PCR duplicates and uniquely aligned reads (alignment records without "XS" tag) were used for downstream analyses. The RPKM values for each 100-bp bin were calculated following the formula “read counts/((bin_length/1000) × (total_reads/10^6)). Peak calling was done by MACS2 with parameters: --nolambda --nomodel.

**Global Proteome and Ubiquitylome Profiling**

Lysis and digest: Samples were lysed in 500 µL SDS lysis buffer (5% SDS, 50 mM TEAB pH 8.5, 2 mM MgCl2, 2 µg/mL Aprotinin, 10 µg/mL Leupeptin, 1 mM PMSF, 50 µM PR-619 (Lifesensors, SI9619), 1 mM Chloroacetamide), disrupted by gentle vortexing, and incubated at room temperature for 15 min. Next, 3 µL of benzonase (250 units/µL) (Thomas Scientific, E1014-25KU) was then added to shear DNA in each sample, followed by mixing and incubation for 15 min at room temperature. Sample lysates were centrifuged for 10 min at 20,000 × g to clear, and the supernatant was transferred for S-Trap digestion. The BCA protein assay was used to estimate protein concentration in the supernatant. Disulfide bonds were reduced in 5 mM DTT for 1 hr at 25°C and 1000 RPM shaking, followed by alkylation of cysteine residues in 10 mM IAA in the dark for 45 min at 25°C and 1000 RPM shaking. The samples were acidified at 1:10 12% phosphoric acid:lysate volume:volume ratio. Proteins in samples were precipitated with 6x sample volume of ice-cold S-Trap buffer (90% methanol, 100 mM TEAB). The buffer with precipitate was thoroughly mixed before being transferred to a S-Trap Midi (Protifi) and then centrifuged at 4000 × g for 1 min. The precipitated proteins were washed 4 times, each with 3 mL S-Trap buffer, and centrifuged at 4000 × g for 1 min. The proteins deposited on the S-Trap were then digested overnight with 350 µL digestion buffer (50 mM TEAB with trypsin and LysC, each at 1:50 enzyme:substrate weight:weight ratio). Specifically, the digestion buffer was passed through each S-Trap column with 1 min centrifugation at 4000 × g and then added back atop of the S-Trap columns. The cartridges were left capped overnight at 25°C. After overnight digestion, peptides were eluted from the S-Trap, first with 500 µL 50 mM TEAB and next with 500 µL 0.1% FA, each for 30 seconds at 1000 × g. The final elution of 500 µL 50% ACN/0.1% FA was centrifuged for 1 min at 4000 × g to clear the cartridge. Eluates were snap frozen and vacuum-centrifuged until dry. Dry peptides were then reconstituted in 30% ACN/0.1% FA, and concentration was estimated using the BCA assay.

Enrichment of K-ε-GG peptides: K-ε-GG peptide enrichment was performed using the UbiFast method as previously described[9,10]. 500 µg peptides from each sample were reconstituted in 250 µL HS bind buffer (Cell Signaling Technology) with 0.01% CHAPS. Next, 5 µL of PBS-washed HS anti-K-ε-GG antibody beads (Cell Signaling Technology, 59322) were added into each peptide sample on a 96-well KingFisher plate (Thermo Fisher Scientific). The plate containing peptides and anti-K-ε-GG antibody beads was covered with foil and incubated for 1 hr at 4°C with end-over-end rotation. After incubation, the foil was removed, and the plate was then processed on the KingFisher Flex as previously described[10]. Briefly, enriched peptides bound to beads were washed, first with 50% ACN/50% HS wash and next with PBS. On-bead TMT labeling of K-ε-GG peptides were done with freshly prepared 400 µg TMTpro reagents (Thermo Fisher Scientific) in 100 mM HEPES for 20 min. The reaction was then quenched with 2% hydroxylamine. Next, beads were washed with HS wash buffer before being released into 100 µL PBS. All sample wells were pooled, and the supernatant was removed. K-ε-GG peptides were eluted from the beads with 2 × 10 min incubation in 100 µL of 0.15% TFA. The eluate was finally desalted using C18 StageTips, snap-frozen, and dried in a vacuum centrifuge.

TMT labeling of K-ε-GG enrichment flow-through for serial proteome analysis: Non-TMT-labeled flow-throughs from K-ε-GG-enrichment were processed for proteome analysis as previously described[11]. Briefly, sample peptides were reconstituted in 1 mL of 1% FA and desalted with 50 mg tC18 SepPak cartridges. Eluted peptides were snap-frozen and vacuum-centrifuged to dry completely. Dried peptides from each sample were reconstituted in 30% ACN/0.1% FA, and their concentration was estimated using the BCA assay. Next, 100 µg peptides from each sample were aliquoted and dried for TMT labeling. Peptides were reconstituted in 20 µL 50 mM HEPES, labeled with 200 µg TMTPro18 reagents at a final concentration of 20% ACN, and incubated for 1 hr at 25°C and 1000 RPM shaking. Labeling reactions were diluted to 2 mg/mL with 50 mM HEPES. After confirming complete labeling and balancing of input material, the TMT labeling reaction was quenched with 3 µL 5% hydroxylamine for 15 min at 25°C and 1000 RPM shaking. All labeled peptide samples were combined, snap-frozen, and vacuum dried. The peptides were then reconstituted with 1 mL 1% FA and desalted with a 100 mg tC18 SepPak. The eluate was snap-frozen and dried to completion in a vacuum centrifuge.

Offline bRP fractionation was conducted to separate peptides over a 96-min gradient with a flow rate of 1 mL/min. Solvent A was 5 mM ammonium formate/2% ACN and solvent B was 5 mM ammonium formate/90% ACN. Next, 96 fractions were concatenated into 24 fractions for proteome analysis, and 5 µg peptides from each fraction were transferred into HPLC vials, frozen, and dried in a vacuum centrifuge. The dried fraction peptides were reconstituted in 3% ACN/0.1% FA, and 1 µg peptides was injected for LC-MS/MS analysis.

<u>LC-MS</u>: Enriched K-ε-GG peptides were reconstituted in 9 µL 3% ACN/0.1% FA, which was injected twice in successive loads of 4 µL onto an Orbitrap Exploris 480 mass spectrometer with FAIMS coupled to a Vanquish Neo UHPLC system (Thermo Fisher Scientific) as previously described[9]. Sample was injected onto a 50°C-heated capillary column (Picofrit with 10 µm tip opening/75 µm diameter, New Objective, PF360-75-10-N-5) packed in-house with approximately 30 cm C18 silica material (1.5 µm ReproSil-Pur C18, Dr. Maisch GmbH). Peptides were separated at a flow rate of 200 nL/min with a linear 154 min gradient from 1.8% solvent B (acetonitrile, 0.1% formic acid), 2 min 5.4% B, 122 min 31.5% B, 130 min 54% B, 133 min 72% B, 144 min 45% B, 149 min 45% B. MS1 spectra were measured with resolution of 60,000, an AGC target of 100%, and a mass range from 350 to 1800 m/z. Up to 10 MS2 spectra per duty cycle were triggered at a resolution of 45,000, an AGC target of 50%, an isolation window of 0.7 m/z, and a normalized collision energy of 32. The FAIMS device was operated in standard resolution mode utilizing the compensation voltages (CVs) of –40, –60, and –80 for the first injection and the CVs of –45, –50, and –70 for the second injection.

All 24 proteome fractions from bRP fractionation were reconstituted in 3% ACN/0.1% FA. For LC-MS/MS analysis, 1 µg peptides from each fraction was injected for LC-MS/MS analysis onto an Orbitrap Exploris 480 mass spectrometer coupled to a Vanquish Neo UHPLC system (Thermo Fisher Scientific) as previously described[9]. The sample was injected onto a capillary column (Picofrit with 10 µm tip opening/75 µm diameter, New Objective, PF360-75-10-N-5) packed in-house with approximately 30 cm C18 silica material (1.5 µm ReproSil-Pur C18, Dr. Maisch GmbH) and heated to 50 °C. Peptides were eluted into the Orbitrap Exploris 480 at a flow rate of 200 nL/min. Each bRP fraction was run on a 110min-method, including a linear 84 min gradient from 94.6% solvent A (0.1% FA) to 27% solvent B (99.9% ACN, 0.1% FA), followed by a linear 9 min gradient from 27% solvent B to 54% solvent B. Mass spectrometry was done in a data-dependent acquisition mode. MS1 spectra were measured with a resolution of 60,000, a normalized AGC target of 300%, and a mass range from 350 to 1800 m/z. MS2 spectra were acquired for the top 20 most abundant ions per cycle at a resolution of 45,000, an AGC target of 30%, an isolation window of 0.7 m/z, and a normalized collision energy of 34. The dynamic exclusion time was set to 20 s, and the peptide match and isotope exclusion functions were enabled.

Data analysis: Mass spectrometry data were processed using Spectrum Mill Rev BI.07.11.216 (proteomics.broadinstitute.org). Raw file extraction retained spectra within a precursor mass range of 600 to 6000 Da and a minimum MS1 signal-to-noise ratio of 25. MS1 spectra within a retention time range of ±45 seconds or within a precursor m/z tolerance of ±1.4 m/z were merged. MS/MS search was performed against a human UniProt database. Digestion parameters were set to "trypsin allow P" with an allowance of 4 missed cleavages. The K-ε-GG MS/MS search included fixed modifications, carbamidomethylation on cysteine and TMTPro on the N-terminus and internal lysine, and variable modifications, acetylation of the protein N-terminus, oxidation of methionine, and K-ε-GG on tryptic peptide – "Ubiquitin Residual GG from Tryp Cut on K". The proteome MS/MS search included fixed modifications, carbamidomethylation on cysteine and TMTPro on the N-terminus and internal lysine, and variable modifications, acetylation of the protein N-terminus, oxidation of methionine, N-term deamidation, and N-term Q-pyroglutamate formation. Restrictions for matching included a minimum matched peak intensity of 40% for K-ε-GG and 30% for proteome, and a precursor and product mass tolerance of ±20 ppm. Validation of peptide-spectrum matches were done using a maximum FDR threshold of 1.2% for precursor charge ranging from 2 to 6. PSMs were further filtered for the proteome in protein polishing autovalidation with the target protein score of 0. TMTpro reporter ion intensities were corrected for isotopic impurities using the afRICA correction method in the Spectrum Mill protein/peptide summary module, which utilizes determinant calculations according to Cramer's Rule.

Protein and K-ε-GG peptide quantification and statistical analysis were performed using the Proteomics Toolset for Integrative Data Analysis (Protigy, v1.0.7, Broad Institute, https://github.com/broadinstitute/protigy). Each K-ε-GG peptide or protein was associated with a log2-transformed expression ratio for every sample condition over the median of all sample conditions. Median-MAD normalization was conducted separately on the K-ε-GG peptide data and the global proteome data. K-ε-GG peptide data were then normalized to the global proteome data using the "panoply_ptm_normalization" module of PANOPLY (PANOPLY, Broad Institute, https://github.com/broadinstitute/PANOPLY/wiki). All K-ε-GG peptide log-ratios in all samples are regressed against the log-ratios of cognate proteins. Resulting residuals are the protein-level normalized K-ε-GG peptide values. After normalization, an empirical Bayes-moderated t test was used to compare treatment groups, using the limma R package. Reported p-values associated with every modified peptide or protein were adjusted using the Benjamini–Hochberg FDR approach.

**IP-MS**

Cell culture: MIA PaCa-2 cells were transduced with the *USP48* CDS (*WT_IsoT2* or *WT_IsoT7*) with a C-terminal V5 tag in pLX_203 or empty vector, selected with puromycin (1 µg/mL), and expanded for 7 days with puromycin maintained in the media. Protein expression of the respective short and long USP48 isoforms was confirmed by probing for V5 or USP48 by western blot.

IP: For each IP, cells were washed twice with ice-cold PBS, pelleted by centrifugation, and lysed with IP Lysis Buffer (Thermo Fisher, 87787) supplemented with Halt Protease Inhibitor Cocktail (Thermo Fisher, 78430) and benzonase (Millipore, 71206) for 30 min on ice with intermittent vortexing. The lysates were centrifuged for 15 min at 13,000 RPM, the supernatant was transferred to a fresh tube, and protein was quantified using the Pierce BCA Protein Assay Kit (Thermo Fisher, A55864). For each IP, 1 mg of input protein was mixed with 50 µL of Anti-V5-tag mAb-Magnetic Beads (MBL International, 103268-788), and then incubated overnight at 4°C with gentle rotation to enable immune complex formation. The next day, beads were pelleted by centrifugation at 300 × g for 5 min at 4°C, washed 5 times with Pierce IP Lysis Buffer, and transferred to the Broad Institute Proteomics Platform.

<u>Preparation of IP/MS samples</u>: IP samples collected and enriched with magnetic beads were washed twice with 200 µL of 50 mM Tris-HCl buffer (pH 7.5), transferred to new 1.5 mL tubes (Eppendorf), and washed 2 more times with 200 µL of 50 mM Tris (pH 7.5) buffer. Samples were incubated in 80 µL of digestion buffer (2 M urea/50 mM Tris, 1 mM DTT, and 5 µg/mL trypsin) for 1 hr at room temperature while shaking at 1000 RPM. Following pre-digestion, 80 µL of each supernatant was transferred into new tubes. Beads were then incubated in 80 µL of the same digestion buffer for 30 min while shaking at 1000 RPM. Supernatant was transferred to the tube containing the previous elution. Beads were washed twice with 60 µL of 2 M urea/50 mM Tris buffer, and these washes were combined with the supernatant. To remove residual beads, the eluates were spun down at 5000 x g for 30 seconds, and the supernatant was transferred to a new tube. Samples were reduced with 4 mM DTT for 30 min at room temperature with shaking. Following reduction, samples were alkylated with 10 mM iodoacetamide for 45 min in the dark at room temperature. An additional 0.5 µg of trypsin was added, and samples were digested overnight at room temperature while shaking at 700 RPM.

Following overnight digestion, samples were acidified (pH<3) with neat formic acid to a final concentration of 1% FA. Samples were then desalted using in-house packed C18 (3M) StageTips. Briefly, C18 StageTips were conditioned sequentially with 100 µL of 100% MeCN acetonitrile, 100 µL of 50% MeCN with 0.1% FA, and 2x 100 µL of 0.1% FA. Next, acidified peptides were loaded onto the C18 StageTips and washed twice with 100 µL of 0.1% FA. Desalted peptides were then eluted from the C18 resin using 50 µL of 50% MeCN/0.1% FA, snap-frozen, and vacuum-centrifuged until completely dry.

Desalted peptides were labeled with TMTpro reagents (Thermo Fisher Scientific). Peptides were resuspended in 80 µL of 50 mM HEPES and labeled with 20 µL 25 µg/µL TMTpro reagents in acetonitrile. Samples were incubated at room temperature for 1 hr with shaking at 1000 RPM. The TMTpro reaction was quenched with 4 µL of 5% hydroxylamine at room temperature for 15 min with shaking. TMTpro-labeled samples were combined, dried to completion, reconstituted in 100 µL of 1.0% FA, and desalted on StageTips. TMTpro labeled peptide sample was fractionated by bRP fractionation. StageTips packed with 2 disks of SDB-RPS (Empore) material. StageTips were conditioned with 100 µL of 100% MeOH, followed by 100 µL of 50% MeCN/0.1% FA and 2 washes with 100 µL of 0.1% FA. Peptide samples were resuspended in 300 µL of 0.1% FA (pH<3) and loaded onto StageTips. Eight stepwise elutions were carried out in 100 µL of 20 mM ammonium formate buffer with increasing concentration: 5%, 7.5%, 10%, 12.5%, 15%, 20%, 25%, and 45% MeCN. Eluted fractions were dried to completion, reconstituted in 9 µL of 3% MeCN/0.1% FA with 4 µL of each fraction injected for MS/MS analysis.

<u>LC-MS analysis of IP samples</u>: Peptide samples were separated with an online nanoflow Proxeon EASY-nLC 1200 UHPLC system (Thermo Fisher Scientific) and run on an Orbitrap Exploris 480 mass spectrometer (Thermo Fisher Scientific). Each sample was injected onto an in-house packed 27 cm x 75 µm internal diameter C18 silica picofrit capillary column (1.9 mm ReproSil-Pur C18-AQ beads, Dr. Maisch

GmbH, r119.aq; PicoFfrit 10 um tip opening, New Objective, PF360-75-10-N-5). The column was heated to 50°C using a column heater sleeve (Phoenix-ST). The mobile phase flow rate was 200 nL/min, comprised of 0.1% formic acid (Solvent A) and 100% acetonitrile/0.1% formic acid (Solvent B). Fractions were analyzed using a 110-min LC– MS/MS method used the following gradient profile: (min:%B) 0:1.8;1:5.4; 85:27; 94:54; 95:81; 100:81; 100.1:45; 109.1:45. Data acquisition was done in the data-dependent mode acquiring HCD MS/MS scans (r = 45,000) after each MS1 scan (r = 60,000) on the top 20 most abundant ions using a normalized MS1 AGC target of 300% and an MS2 AGC target of 30%. The maximum ion time utilized for MS/MS scans was 105 ms; the HCD-normalized collision energy was set to 34; the dynamic exclusion time was set to 20 seconds, and the peptide match and isotope exclusion functions were enabled. Charge exclusion was enabled for charge states that were unassigned, 1 and >6.

<u>Data analysis</u>: Data was processed using Spectrum Mill (proteomics.broadinstitute.org). For all samples, extraction of raw files retained spectra within a precursor mass range of 750-6000 Da and a minimum MS1 signal-to-noise ratio of 25. MS1 spectra within a retention time range of +/- 45 seconds, or within a precursor m/z tolerance of +/- 1.4 m/z were merged. MS/MS searching was performed against a human Uniprot database (release date 09/02/2021) with manually inputted annotations for USP48 WT_IsoT2 and WT_IsoT7 constructs. Digestion parameters were set to ‘‘trypsin allow P’’ with an allowance of 4 missed cleavages. The MS/MS search included fixed modification of carbamidomethylation on cysteine. TMTpro was searched using the full-mix function. Variable modifications were acetylation, oxidation of methionine, pyroglutamic acid, and deamidated amines. Restrictions for matching included a minimum matched peak intensity of 30% and a precursor and product mass tolerance of +/- 20 ppm. Peptide spectrum matches (PSMs) were validated using a maximum false discovery rate (FDR) threshold of 0.8% for a precursor charge range of 2-4, and 0.4% for a range of 5-6. Protein level autovalidation was performed to a target protein-level FDR of 0. A subgroup top (SGT) protein/protein report was generated, and TMTpro reporter ion intensities were corrected for isotopic impurities in the Spectrum Mill protein/peptide summary module using the afRICA correction method, which implements determinant calculations according to Cramer's Rule.

To correctly identify the USP48 protein isoforms derived from WT_IsoT2 and WT_IsoT7, unique protein scores (uniqueScores) were generated for each protein in a protein group. This score is calculated by summing the identification score of each peptide used to identify a unique protein. The top-scoring protein is then used as the identified protein for downstream analysis.

For statistical analysis, log2 multi-median reporter ion ratios were used. Reporter ion ratios were median-MAD normalized, filtered for up 70% missing values, and a two-sample moderated t test was performed. Adjusted p-value were calculated using the Benjamini–Hochberg FDR approach, and identified proteins with an adjusted p-value<0.05 were considered significant.

**USP48 CDS Sequences**

The following *USP48* CDS sequences were used for rescue and conditional degradation experiments. The C98S mutation was introduced into the *WT_IsoT7* CDS via site-directed mutagenesis by Genscript to generate the *C98S_IsoT7* CDS.

<u>USP48_IsoT2 (short isoform) with V5 tag (codon optimized)</u>

ATGGCCCCTAGACTGCAACTGGAAAAAGCCGCTTGGCGGTGGGCCGAAACCGTGCGACCTG
AGGAGGTGTCTCAGGAGCATATTGAGACCGCCTATAGAATCTGGCTGGAACCTTGTATCAGA
GGCGTGTGCAGACGGAACTGCAAGGGAAATCCTAATTGCCTGGTCGGCATCGGCGAGCACA
TCTGGCTGGGCGAAATCGACGAGAACTCCTTCCACAACATCGATGATCCCAACTGCGAGAGA
AGAAAAAAGAACAGCTTCGTGGGCTTAACCAACCTGGGCGCTACATGCTACGTTAACACCTT
CCTGCAGGTGTGGTTCCTGAATCTGGAACTGAGACAGGCCCTGTACCTGTGCCCTTCTACCTG
TTCTGATTACATGCTGGGCGACGGCATCCAGGAGGAAAGGACTACGAGCCTCAGACCATCT
GCGAGCACCTGCAGTACCTGTTTGCCCTGCTGCAGAACAGCAACAGACGGTACATCGACCCT
AGCGGATTTGTGAAGGCCCTGGGACTTGACACAGGCCAGCAGCAGGACGCTCAGGAGTTCA
GCAAGCTGTTTATGAGCCTGCTCGAGGACACCCTGTCCAAGCAGAAGAACCCTGATGTGCGG
AACATCGTGCAGCAACAATTTTGTGGCGAGTACGCCTACGTGACCGTGTGCAACCAGTGCGG
AAGAGAGAGCAAACTGCTGTCCAAGTTCTACGAGCTGGAACTCAACATCCAGGGCCACAAG
CAGCTGACCGACTGCATCAGCGAATTCCTCAAGGAAGAAAAGCTGGAAGGCGATAACAGAT
ACTTCTGCGAAAACTGTCAGTCTAAACAAAATGCCACACGGAAGATCCGGCTGCTGAGCCTG
CCTTGCACCCTGAACCTGCAGCTGATGAGGTTCGTGTTCGACAGACAGACCGGCCACAAGAA
GAAGCTGAACACATACATCGGCTTCAGCGAGATCCTGGACATGGAACCATACGTGGAACAT
AAAGGCGGAAGCTACGTGTACGAGCTGTCTGCCGTGCTGATCCACAGAGGCGTGTCCGCCTA
TTCTGGCCACTACATCGCCCACGTGAAGGACCCCCAGAGCGGCGAATGGTATAAGTTCAACG
ACGAGGACATCGAAAAGATGGAAGGCAAGAAACTGCAGCTGGGCATCGAGGAGGATCTGG
CCGAGCCAAGCAAGAGCCAGACCAGAAAGCCCAAGTGCGGCAAAGGTACACACTGTAGTAG
AAATGCCTACATGCTGGTGTACCGCCTGCAAACACAGGAGAAGCCTAACACCACAGTCCAG
GTGCCCGCCTTCCTGCAGGAGCTGGTGGACCGGGACAATAGCAAATTCGAGGAATGGTGTAT
TGAAATGGCTGAGATGCGGAAGCAGAGCGTGGACAAGGGCAAGGCCAAGCACGAGGAGGT
GAAAGAGCTGTACCAGAGACTGCCTGCTGGCGCCGGACTGGGCAAGCCTATCCCCAACCCC
CTGCTCGGCCTGGATAGCACATGA

USP48_IsoT7 (long isoform) with V5 tag (codon optimized)
ATGGCCCCTAGACTGCAGCTCGAGAAGGCTGCCTGGAGATGGGCCGAGACAGTGCGGCCTG
AAGAAGTGAGCCAGGAGCATATCGAGACAGCCTATAGAATCTGGCTCGAACCCTGCATCCG
GGGCGTCTGCAGAAGGAACTGCAAGGGAAATCCTAACTGTCTGGTGGGCATCGGCGAGCAC
ATCTGGCTGGGAGAAATCGACGAGAATAGCTTTCACAACATCGACGACCCTAACTGTGAAA
GAAGGAAAAAAAATAGCTTCGTGGGACTGACCAACCTGGGCGCCACCTGCTACGTGAACAC
ATTCCTGCAAGTCTGGTTCCTGAATCTGGAGCTAAGACAGGCTCTGTACCTGTGCCCTAGCA
CCTGTTCTGATTACATGCTGGGGGACGGCATCCAGGAGGAAAGGATTATGAGCCACAGAC
CATCTGCGAGCACCTGCAATACCTGTTCGCCCTGCTGCAAAACAGCAACAGAAGATACATCG
ATCCTAGCGGCTTTGTAAAGGCCCTGGGCCTGGACACAGGCCAGCAGCAGGACGCCCAGGA
GTTCAGCAAGCTGTTCATGAGCCTGTTGGAAGATACACTGTCCAAACAAAAGAACCCTGACG
TGCGGAACATCGTGCAGCAGCAATTTTGCGGCGAGTACGCCTACGTGACAGTGTGCAACCAG
TGCGGCAGAGAGAGCAAGCTGCTGAGCAAGTTCTACGAGCTGGAGCTGAACATTCAGGGCC
ACAAGCAGCTGACTGATTGCATAAGCGAGTTCCTGAAGGAAGAAAAGCTGGAAGGCGACAA
CAGATACTTCTGCGAGAACTGCCAATCCAAGCAGAACGCCACTAGAAAAATCCGGCTGCTG
AGCCTGCCTTGCACCCTGAATCTGCAGCTCATGCGGTTCGTGTTCGATAGACAGACCGGCCA
CAAGAAGAAGCTGAACACCTACATTGGCTTCAGCGAGATCCTGGATATGGAACCTTACGTGG
AACACAAAGGCGGCAGCTACGTGTACGAGCTGTCAGCCGTGCTGATCCACCGGGGCGTGAG
CGCCTACTCCGGCCACTACATCGCCCACGTGAAGGACCCCCAGAGCGGCGAATGGTACAAG

TTCAACGACGAGGACATCGAGAAGATGGAAGGCAAGAAGCTGCAGCTGGGTATCGAGGAG
GACCTGGCTGAACCCAGCAAGAGCCAGACCAGAAAGCCTAAGTGCGGAAAGGGCACCCACT
GCTCGAGAAACGCCTACATGCTGGTTTACAGACTGCAGACCCAGGAGAAACCTAACACCAC
CGTGCAAGTGCCCGCCTTTCTGCAGGAGCTCGTGGACAGAGATAATAGCAAGTTCGAGGAAT
GGTGCATCGAAATGGCCGAGATGAGAAAGCAGAGCGTCGACAAGGGCAAAGCTAAGCACG
AGGAGGTGAAGGAGCTGTACCAGAGACTGCCAGCCGGCGCCGAACCCTACGAGTTTGTGTC
TCTGGAATGGCTGCAGAAGTGGCTGGATGAAAGCACCCCTACCAAGCCAATTGATAATCAC
GCTTGCCTCTGTAGCCACGACAAGCTGCATCCTGATAAGATCAGCATCATGAAGCGGATCTC
TGAGTACGCCGCTGATATCTTCTACTCCAGATACGGCGGAGGACCTAGGCTGACCGTGAAGG
CACTCTGCAAAGAGTGCGTGGTGGAAAGATGCAGAATCCTGAGACTGAAGAACCAGCTGAA
CGAGGACTACAAGACAGTGAACAACCTGCTGAAGGCCGCCGTGAAAGGCGACGGCTTCTGG
GTCGGCAAGAGCTCTCTGCGCAGCTGGCGGCAGCTGGCCCTGGAGCAGCTGGACGAACAGG
ATGGCGACGCCGAACAGTCTAATGGCAAGATGAACGGCTCTACCCTGAACAAGGACGAGAG
CAAGGAGGAACGGAAGGAGGAAGAAGAGCTGAATTTCAACGAGGACATCCTGTGTCCTCAT
GGAGAGCTATGTATCAGCGAAAACGAGCGGCGGCTTGTGTCCAAAGAGGCTTGGTCCAAGC
TGCAGCAGTACTTTCCAAAGGCCCCTGAATTCCCCAGCTACAAGGAGTGTTGTTCTCAGTGT
AAAATCCTGGAAAGAGAGGGCGAGGAAAACGAGGCCCTGCACAAGATGATCGCAAACGAA
CAGAAAACGAGCCTGCCTAACCTGTTCCAGGACAAGAACAGACCTTGTCTGAGCAACTGGC
CCGAGGACACCGACGTGCTGTACATCGTGTCCCAGTTCTTCGTGGAAGAGTGGCGGAAATTC
GTTAGAAAGCCCACCCGGTGCTCCCCTGTGAGCAGCGTGGGCAACTCTGCCCTGCTGTGCCC
CCACGGCGGCCTGATGTTCACCTTCGCCAGCATGACCAAAGAGGACAGCAAGCTAATTGCCC
TGATCTGGCCTTCTGAATGGCAGATGATCCAGAAGCTGTTTGTCGTGGACCACGTGATCAAG
ATCACCCGGATCGAGGTGGGCGATGTGAACCCTAGTGAGACACAGTATATCAGCGAACCCA
AACTGTGCCCCGAATGTCGCGAGGGACTGCTTTGCCAGCAGCAACGGGACCTGAGAGAGTA
CACCCAAGCTACCATCTACGTGCACAAGGTGGTGGACAACAAGAAGGTGATGAAAGATAGC
GCCCCTGAGCTGAACGTGTCTAGCAGCGAGACGGAAGAGGACAAAGAGGAAGCCAAGCCA
GACGGCGAGAAGGACCCCGACTTCAATCAGTCTAACGGCGGCACCAAGAGACAGAAAATCT
CTCACCAGAACTACATCGCCTACCAAAAGCAGGTGATCAGAAGAAGTATGCGCCACAGAAA
GGTTCGAGGCGAGAAGGCCCTGCTGGTGAGCGCTAATCAGACACTGAAGGAGCTGAAGATC
CAGATCATGCACGCCTTCTCGGTCGCCCCCTTTGACCAGAACCTGAGCATCGACGGCAAAAT
TCTGAGCGATGACTGCGCCACACTGGGCACCCTGGGCGTGATCCCTGAGAGCGTGATCCTGC
TGAAGGCCGATGAACCTATCGCCGACTACGCCGCCATGGACGATGTGATGCAAGTGTGCATG
CCTGAGGAGGGCTTTAAGGGAACCGGCCTGCTGGGACACGGCAAGCCTATCCCCAATCCCCT
GCTGGGACTGGACAGCACATGA

**Vectors and sgRNA Sequences**

Individual sgRNAs were cloned into expression vectors carrying the *PAC* (puromycin resistance), *mCherry*, *GFP*, *BFP*, or *RFP657* CDS: pXPR_003 (Broad GPP), pXPR_050 (Broad GPP), pXPR_044 (Broad GPP), pLentiGuide-Puro (Addgene, 52963), or pLentiGuide modified to have a fluorophor CDS (*GFP*, *BFP*, or *RFP657*) followed by a P2A sequence and *PAC* cassette (Genscript). The following sgRNA sequences were used in this study.

| | *Gene* | *Broad ID* | *Target Sequence* |
|---|---|---|---|
| sgCTRL_20 (sg1) | Ctrl (LacZ) | BRDN0001058320 | AACGGCGGATTGACCGTAAT |

| | | | |
|---|---|---|---|
| sgCTRL_35 (sg2) | Ctrl (Luciferase) | BRDN0000585635 | AATTGTCTTGTCCCTATCGA |
| sgCTRL_00 (sg3) | Ctrl (Intergenic) | BRDN0004438500 | TTTCGGACCACGGTCGACCA |
| sgUSP48_06 (sg1) | *USP48* | BRDN0003512706 | AGACTGTGGATCTTTCACGT |
| sgUSP48_80 (sg2) | *USP48* | BRDN0003827980 | TACTACACATTCCTTACACA |
| sgUSP48_63 (sg3) | *USP48* | BRDN0003514063 | TGAGATGCGTAAGCAAAGTG |
| sgEZH2_51 (sg1) | *EZH2* | BRDN0000733051 | TTATGATGGGAAAGTACACG |
| sgEZH2_85 (sg2) | *EZH2* | BRDN0003228385 | AATAATTGCACTTACGATGT |
| sgBCOR_22 (sg1) | *BCOR* | BRDN0001493922 | GTGCAGACTGGAGAATACAG |
| sgBCOR_69 (sg2) | *BCOR* | BRDN0001493969 | AGTTAGCAGCGAGTTCCCCG |
| sgDCK_59 (sg1) | *DCK* | BRDN0001148859 | TCAAGAAAATCTCCATCGAA |
| sgDCK_71 (sg2) | *DCK* | BRDN0001145971 | TGTATGAGAAACCTGAACGA |
| sgUCK2_52 (sg1) | *UCK2* | BRDN0001146352 | TCTGCTCCGAGGTAAGGACA |
| sgUCK2_69 (sg2) | *UCK2* | BRDN0001145769 | CTTCCTTATAGGCGTCAGCG |
