## Supplementary Table Legends for "Discovery of chromatin-based determinants of azacytidine and decitabine anti-cancer activity"

### SUPPLEMENTARY TABLES

**Supplementary Table 1:** 6-day PRISM screen viability results. List of L2FC and AUC values for each PRISM cell line treated with AZA, DEC, or GSK-3685032 for 6 days.

**Supplementary Table 2:** 6-day PRISM screen RNA and protein biomarker results. Effect sizes and p-values are shown for protein and RNA expression features for the AZA (0.25 μM) and DEC (0.5 μM) PRISM screens.

**Supplementary Table 3:** Proteome and ubiquitylome results. Paired proteome and ubiquitylome profiling was performed for MOLM-13, MV4;11, and MIA PaCa-2 following USP48 knockout or GSK-3685032 treatment.

**Supplementary Table 4:** Humagne CRISPR/Cas12a drug modifier screen results. List of gene-level L2FC values, Z-scores, and p-values for the AZA and GSK-3685032 modifier screens in MOLM-13, MV4;11, and U-937.

**Supplementary Table 5:** GSEA results. Top reactome gene sets represented by the 37 sensitizer gene hits identified in the Humagne drug modifier screens.

**Supplementary Table 6:** Bison CRISPR/Cas9 drug modifier screen results. List of gene-level enrichment scores and p-values for the DEC and GSK-3685032 modifier screens in KARPAS-299 and MIA PaCa-2.

**Supplementary Table 7:** CRISPR/Cas9 PRISM screen viability results. List of L2FC for each cell line in the *USP48*-intact and *USP48*-knockout pools treated with GSK-3685032.

**Supplementary Table 8:** ChIP-seq results. Peak calls for the USP48 ChIP-seq experiments in MOLM-13.

**Supplementary Table 9:** ATAC-seq results. Peak calls for the ATAC-seq experiments in MOLM-13.

**Supplementary Table 10:** IP-MS interactome results for the short USP48 isoform (T2) and long USP48 isoform (T7).
